# MLL family members regulate H3K4 methylation to ensure CENP-A assembly at human centromeres

**DOI:** 10.1101/2022.06.20.496844

**Authors:** Kausika Kumar Malik, Sreerama Chaitanya Sridhara, Kaisar Ahmad Lone, Payal Deepakbhai Katariya, Shweta Tyagi

**Author notes:** Corresponding author: Laboratory of Cell Cycle Regulation, CDFD Uppal, Hyderabad 500039, India.

## Abstract

The active state of centromeres is epigenetically defined by the presence of CENP-A interspersed with histone H3 nucleosomes. While the importance of dimethylation of H3K4 mark for centromeric transcription has been highlighted in various studies, the identity of the enzyme(s) depositing these marks on the centromere is still unknown. The MLL (KMT2) family play a crucial role in RNA polymerase II (Pol II)-mediated gene regulation by methylating H3K4. Here, we report that MLL family regulate transcription of human centromeres. CRISPR-mediated downregulation of MLL causes loss of H3K4me2, resulting in an altered epigenetic chromatin state of the centromeres. Intriguingly, our results reveal that loss of MLL, but not SETD1A, increases co-transcriptional R-loop formation, and Pol II accumulation at the centromeres. Finally we report that the presence of MLL and SETD1A is crucial for kinetochore maintenance. Altogether, our data reveals a novel molecular framework where both the H3K4 methylation mark and the methyltransferases regulate stability and identity of the centromere.

## Introduction

Centromeres are specialized regions on chromosomes that form a scaffold of the kinetochore, a multi-protein complex that links chromosome to spindle microtubules to facilitate faithful chromosome segregation during mitosis; failure of this process leads to chromosomal structural and numerical abnormalities often seen in pathological conditions such as cancer (Thompson and Compton., 2011). Centromeres are organized into two broad regions, the inner ‘core’ centromere region flanked by large outer pericentromere. The centromere is characterized by repetitive α-satellite DNA sequences, which consist of ∼171 bp monomers organized in tandem to form higher-order repeat (HOR) arrays that range from 2 to 5 Mb and are species- and chromosome-specific (Corless et al., 2020). Although, the function of the centromere is highly conserved among the eukaryotes, the α-satellite DNA sequences are not evolutionary conserved (Thakur et al., 2021). In fact, centromeres pose an evolutionary conundrum as they are epigenetically defined by — centromeric protein A (CENP-A) — a Histone 3 variant, and not by the presence of α-satellite DNA. Interestingly, centromere chromatin (here on centrochromatin) constitutes CENP-A nucleosomes interspersed with Histone 3 nucleosomes, bearing post-translational modifications such as histone 3 lysine 4 dimethylation (H3K4me2), lysine 9 acetylation (H3K9ac), and lysine 36 dimethylation (H3K36me2) providing a unique chromatin state (Molina et al., 2016; Allshire and Ekwall, 2015; Bergmann et al., 2011; Ribeiro et al., 2010; Sullivan and Karpen., 2004). Although, initially thought to be transcriptionally silent, centrochromatin is now known to be transcribed by RNA polymerase II (RNA Pol II) and produces centromere RNA (cenRNA) transcripts (Bury et al., 2021; Choi et al., 2011; Quénet and Dalal., 2014; Chan et al., 2012; Bergmann et al., 2011; Wong et al., 2007). Moreover, histone modification, centromere transcription, and the cenRNA are important for the centromere and kinetochore assembly and function (Bobkov et al., 2018; McNulty et al., 2017; Blower, 2016; Molina et al., 2016; Grenfell et al., 2016; Quénet and Dalal, 2014; Ideue., 2014; Fachinetti et al., 2013; Bergmann et al., 2011; Ferri et al., 2009; Wong et al., 2007; Sullivan and Karpen, 2004; Nakano et al., 2003). For instance, cenRNAs physically interacts with CENP-A, centromeric protein C (CENP-C), and Holliday junction recognition protein (HJURP) to efficiently recruit these proteins at the centromeres (McNulty., 2017; Quénet and Dalal., 2014). Furthermore, several reports suggest that RNA Pol II-mediated centromere transcription and cenRNA ensure loading of CENP-A at the centromeres in a cell cycle-specific manner (McNulty et al., 2017; Molina et al., 2016 Quenet and Dalal., 2014; Bergmann et al., 2011; Wong et al., 2007). The specialized nature of centrochromatin has led various groups to investigate the importance of histone modifications and transcription at the centromeres, and kinetochore maintenance (Martins et al., 2016; Molina et al., 2016; Chen et al., 2015; Rosic et al., 2014; Bergmann et al., 2011; Lu and Gilbert, 2007; Sullivan and Karpen, 2004). Using synthetic human artificial chromosome (HAC), Earnshaw’s group has shown that H3K4me2 mark is not only essential for cenRNA transcription, but its removal also resulted in rapid loss of transcription leading to impaired CENP-A incorporation and eventually, kinetochore instability (Molina et al., 2016; Bergmann et al., 2011). While the importance of the H3K4me2 mark for centromeric stability has been revealed, the identity of the histone methyltransferases (HMT) depositing this mark on the centromere remains elusive.

In eukaryotes, the lysine methyltransferase 2 (KMT2, SET1 or MLL or) family of proteins deposit the H3K4 methylation marks. In humans, there are six members in this family including MLL1 (MLL or Mixed lineage leukemia protein), MLL2, MLL3, MLL4, SET Domain Containing 1A (SETD1A), and SETD1B. While SETD1A is a global H3K4 tri-methyltransferase, MLL1-4 displays locus-specific methylation activity (Sugeedha et al., 2020; Crump and Milne, 2019; Rao and Dou., 2015). All members of this family activate transcription through the <u>S</u>u(var)3-9, <u>E</u>nhancer-of-zeste, <u>T</u>rithorax (SET) domain which is responsible for the methyltransferase activity of these enzymes. In addition, some members like MLL and MLL2 also use the transcription activation domain (TAD) to promote transcription (Goto et al., 2002). Different reports implicate MLL family members in the assembly of the transcription pre-initiation complex and recruitment of RNA Pol II to target genes (Smith et al., 2011). In fact, H3K4 methylation has been proposed to be a prerequisite for recruitment of the basal transcription machinery and initiation of transcription for several mammalian gene targets (Wang et al., 2009; Vermeulen et al., 2007). However, how this how this machinery works in the context of an active intergenic chromatin state such as centromeres is still not clear.

MLL family members are involved in a wide variety of roles. However, the role of these proteins in mitosis is recently coming to light. Interestingly, all mitotic functions described so far for the members of this family are involved in averting chromosome mis-segregation and thus maintaining genomic integrity (Ali et al., 2017; Schibler et al., 2016; Ali et al., 2014; Latham et al., 2011; Zhang et al., 2005). We have previously reported the localization of MLL and SETD1A on spindle apparatus, and shown that MLL regulates proper chromosome alignment and segregation using protein-protein interactions (Ali et al., 2017). Here, we show that most MLL family members have a role in regulating transcription of cenRNA. We report that endogenous MLL and SETD1A bind to human centromeres and regulate centromere transcription. Furthermore, using MLL knock-out cell lines, we reveal MLL as the ‘writer’ for H3K4me2, and its crucial role in sustaining the unique epigenetic state of the human centromeres. Interestingly, removal of MLL but not SETD1A augments centromere R-loops (or co-transcriptional RNA:DNA hybrids) and RNA Pol II at centromeres. We also observe that loss of MLL and SETD1A adversely affects kinetochore maintenance as recruitment of CENP-B, CENP-C, and HJURP are compromised. Finally, we show that MLL and SETD1A affect the loading of nascent CENP-A during the early G1 phase. Our results provide insights into centromere transcription and reveal a functional difference between the different members of the MLL family in regulating intergenic transcription.

## Results

### Members of the MLL family regulate centromeric transcription

Several studies have shown that centromeres are transcribed by RNA Pol II in a unique environment on chromatin decorated with histone modification like H3K4me2 and H3K36me2 (Molina et al., 2016; Chen et al., 2015; Bergmann et al., 2011; Sullivan and Karpen, 2004). However, how this transcription is regulated, is not fully understood. As members of the MLL family are responsible for depositing the H3K4me2 marks on the genome, we postulated that they regulate transcription of centromeric RNA (cenRNA). To test this hypothesis, we used previously characterized siRNAs to knock down various members of the MLL family (Ali et al., 2014) and studied the effect on α-satellite cenRNAs. These were detected by quantitative real-time polymerase chain reaction (qRT-PCR) using universal primer set from α-satellite DNA, sequences which are present on all centromeres (Quénet and Dalal., 2014; Dunham et al., 1992). We observed about 50% decrease in MLL family member transcripts, which resulted in a similar decrease in transcription of α-satellite arrays (Figure 1A, S1A). Several studies have reported use of RNA Pol II specific inhibitors reduce cenRNA transcripts (Chen et al., 2021; Bury et al., 2020; Quénet and Dalal., 2014; Wong et al., 2007). We used these Pol II inhibitors as a positive control in our experiments and observed decreased transcription from the centromeres upon treatment with Triptolide, α-amanitin and CDK9 inhibitor—LDC000067 hydrochloride (Figure 1B).

**Figure 1:**
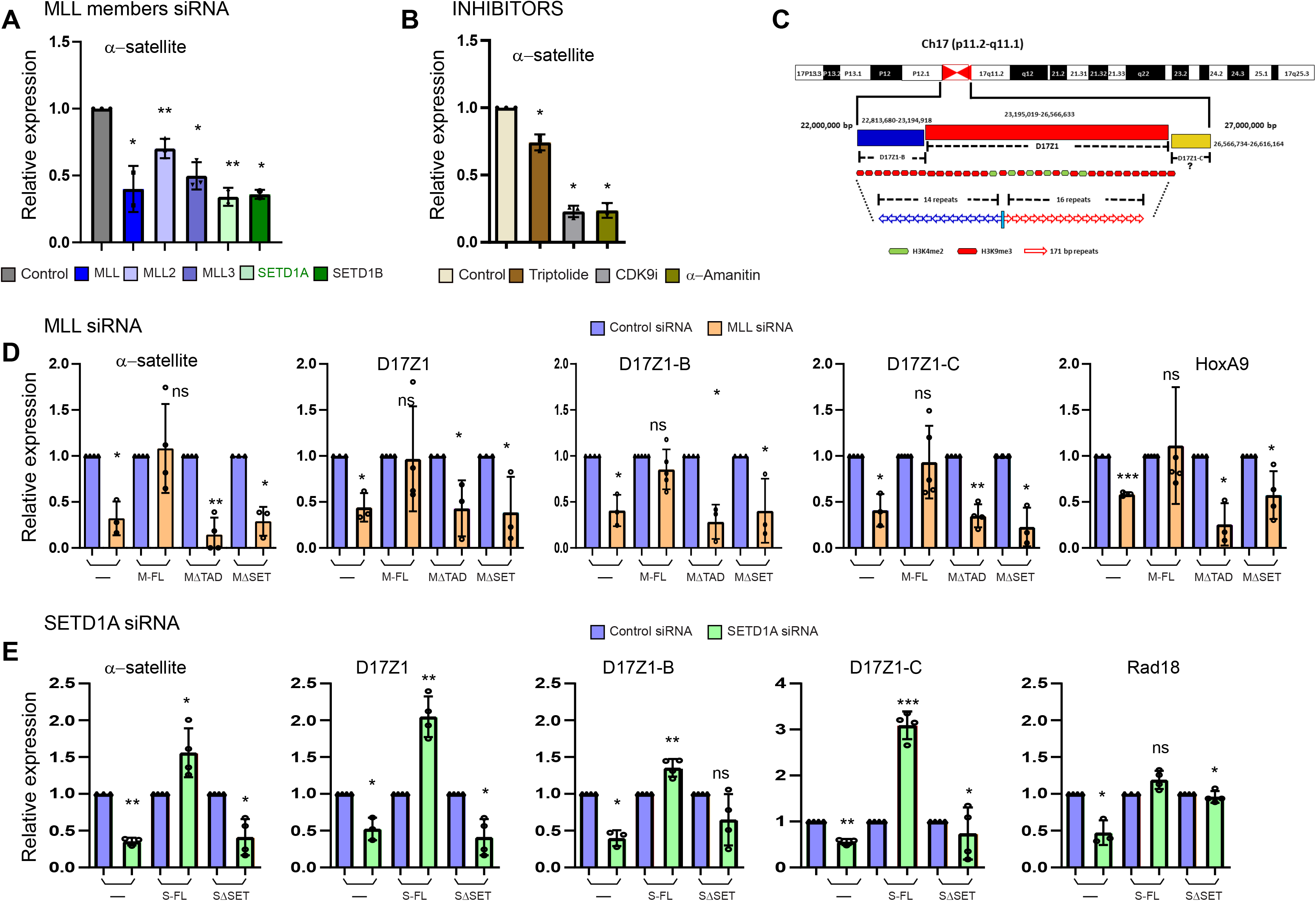
RNAi-mediated downregulation of MLL family members abrogates centromeric transcription. (**A**) Shown is a qRT-PCR analysis of universal α-satellite cenRNA transcript level in Control, MLL, MLL2, MLL3, MLL4, SETD1A, and SETD1B siRNA treated cells as indicated. (**B**) qRT-PCR analysis of α-satellite cenRNA expression after treatment with either Control (DMSO), Triptolide (20 µM), CDK9 inhibitor (20 µM), α-amanitin (20 µg), or for 4 hrs is shown. (**C**) Schematic representation of human chromosome 17 HOR α-satellite arrays—D17Z1, D17Z1-B, and D17Z1-C. The numbers indicate base pairs and are based on GRCh 38/hg38. (**D**) Following Control or MLL siRNA treatment, cenRNA transcript levels of α-satellite arrays as indicated were measured in parent U-2OS cells (—) or U-2OS cells stably expressing full-length MLL (M-FL), TAD deleted MLL (MΔTAD) and SET domain deleted MLL (MΔSET). (**E**) Shown is a qRT-PCR analysis of cenRNA transcripts as indicated, following treatment with Control or SETD1A siRNA in parent U-2OS cells (—) or U-2OS cells expressing full-length SETD1A (S-FL) and SET domain deleted SETD1A (SΔSET). cDNA was synthesized from total RNA after rigorous DNase treatment and amplified using qRT-PCR for indicated RNAs. Data from all samples were normalized to GAPDH mRNA levels from respective samples by using − ΔΔCT method and expression is shown relative to control siRNA-treated/DMSO treated cells from respective cell line/treatment (which is arbitrarily set to 1). Data obtained for the α-satellite transcript in **A** from MLL and SETD1A siRNA treatment of parent U-2OS cells are replotted in **D** and **E** respectively, for ease of comparison. Each experiment was performed at least three, or more times except α-amanitin treatment (two times). Error bars represent SD. *P ≤ 0.05, **P ≤ 0.005, *** P ≤ 0.0005, ns: not significant P > 0.05 (two-tailed Student’s t-test). CDK9i, CDK9 inhibitor.

As α-satellite DNA sequences are also known to be present in peri-centromeres, in addition to the universal α-satellite primer, we used well-characterized primers from chromosome 17 specific α-satellite arrays (Figure 1C; McNulty et al., 2017). Chromosome 17 contains three α-satellite arrays— D17Z1, D17Z1-B and D17Z1-C—which vary in their sequence, and size of their HORs (McNulty and Sullivan., 2018). These arrays are functionally distinct, producing active as well as inactive array transcripts (McNulty et al., 2017). HORs of both D17Z1 and D17Z1-B have been shown to form centromeres independently (Maloney et al., 2012) while the status of D17Z1-C is not clear (Hayden et al., 2013). In addition, we analysed for genes that are co-regulated by RNA Pol II and MLLs like *HOXA9, PAX3*, and *RAD18* (Alsulami et al., 2019; Wang et al., 2009). Once we knocked down different MLL family members, we observed a reduction in all three array-specific transcripts from chromosome 17 (Figure 1D-E, S1B). Treatment with the three Pol II inhibitors, similarly reduced cenRNA transcripts from D17Z1, D17Z1-B, and D17Z1-C (Supplementary Figure S1C). To sum up, our results indicate that all members of the MLL family tested here facilitate RNA Pol II-mediated transcription of cenRNA.

### MLL and SETD1A require the SET domain to regulate centromeric transcription

In order to understand how MLLs regulate cenRNA transcription, we choose MLL and SETD1A to study the process further because loss of both these proteins is known to produce chromosome-segregation defects (Ali et al, 2017, 2014),— defects, which are also caused as a consequence of the perturbed transcription at the centromere (McNulty et al., 2017; Molina et al., 2016; Grenfell et al., 2016; Chen et al., 2015; Quénet and Dalal., 2014; Rosic et al., 2014; Chan et al., 2012; Bergmann et al., 2011). Reduced transcripts or protein levels of MLL and SETD1A resulted in a reduction of both α-satellite as well as array-specific transcripts from chromosome 17 (Figure 1D-E, S1A,D-E). In order to determine if this reduction was specific to MLL and SETD1A, we analysed transcripts when siRNA-treated cells were complemented with full-length protein (Figure 1D-E, S1F-G). We also used SET-domain deleted protein(s) to determine the role of the SET domain in cenRNA transcription. We utilized stable cell-lines made in U-2OS cells for this purpose (Figure S1F, Ali et al., 2014, this study). To ensure that only endogenous *MLL* or *SETD1A* transcript is affected by our siRNA treatment, and not the recombinant one, we made use of siRNA directed against 3′ UTR of precursor mRNA (MLL siRNA #2) or made recombinant constructs siRNA resistant by introducing silent mutations (Figure S1G, SETD1A siRNA#1). Our findings indicate that MLL and SETD1A specifically regulate centromeric transcription and this regulation is dependent on the SET domain. Interestingly, when we studied the TAD deletion in MLL (MLLΔTAD), we found that TAD also affects the transcript levels at the centromere(Figure 1D-E, S1F-G). Taken together, our findings suggest that MLL and SETD1A use their transcription-competent domains to regulate cenRNA transcription from both active as well as inactive arrays.

### MLLs bind to the human centromere repeats

MLL regulate the transcription of a large number of genes, either by direct binding to gene targets or indirectly (Wang et al., 2009; Blobel et al., 2009; Milne et al., 2005; Guenther et al., 2005; Schraets et al., 2003). Therefore, we wanted to investigate if MLL and SETD1A were present on the centromeres to regulate the cenRNA transcription. We checked for the presence of MLLs on the centromere using immunofluorescence staining (IF). Consistent with their role in transcription of cenRNA, we found MLL and SETD1A co-localizing with CENP-A on the centromere in mitosis (Figure 2A). As cenRNA transcription has also been reported in early G1 cells, we performed IF in cells synchronized in this stage (Figure 2B). MLL and SETD1A showed co-localization with CENP-A not only in early G1 but in asynchronous interphase cell population as well (Figure 2B, S2A). In contrast to the mitotic cells and consistent with their regulatory role in the genome, MLL and SETD1A localization on the centromere was distinct but not exclusive (Figure 2A-B, S2A). In order to further validate the specific signal of our proteins on the centromere, we performed siRNA-mediated depletion of MLL or SETD1A and observed reduced staining of these proteins, further confirming that MLL and SETD1A are specifically present on the centromeres (Supplementary Figure S2B-C). This was accompanied by a no primary antibody control (Supplementary Figure S2D). We also checked for the presence of MLL2, MLL3 and SETD1B on the centromere (Supplementary Figure S2E). Even though not as distinct as MLL or SETD1A, may be due to the fixation conditions or antibody used, these proteins were present on the centromeres (Supplementary Figure S2E). Taken together our data suggests that in addition to the canonical non-repetitive ‘regular’ genomic loci, MLLs also bind to, and regulate transcription from repetitive centromeric sequences transcribing non-coding RNA.

**Figure 2.**
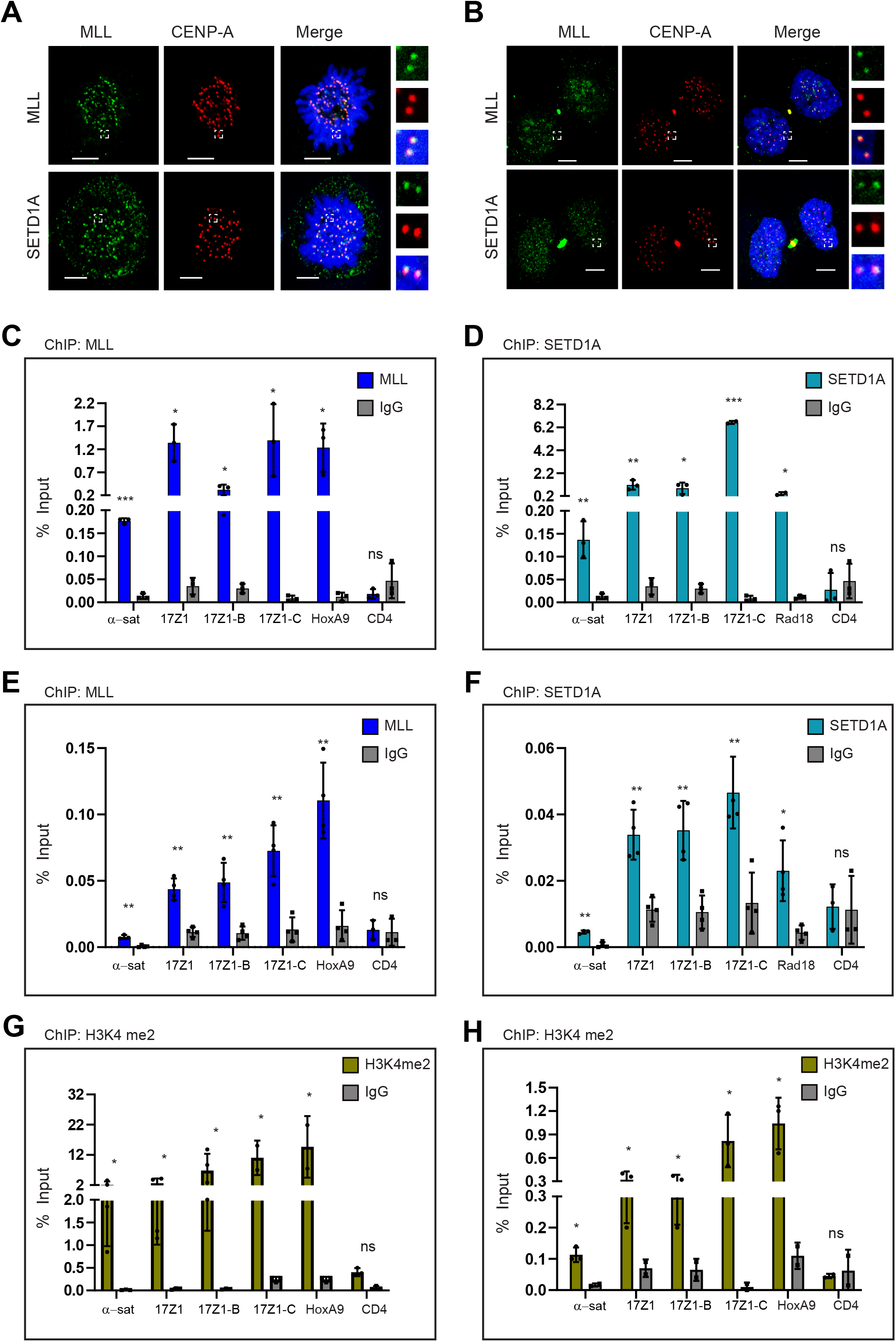
MLLs binds to the human centromere repeats. (**A-B**) Immunofluorescence staining (IF) of endogenous MLL (green) or SETD1A (green) with CENP-A (red) in U-2OS cells synchronized in mitosis (**A**) or early G1 (**B**) is shown. DNA was stained with DAPI (blue). The area in the white square is magnified and shown on the right for each image. Scale bar, 5μm. Pearson correlation coefficient was measured for more than 100 centromeres and mean with SEM is shown between CENP-A and — MLL (**A**)=0.45+0.015, SETD1A(**A**)=0.50±0.014, MLL(**B**)=0.<u>3</u>5+0.015, and SETD1A(**B**) =0.50±0.013. (**C-D**) Chromatin immunoprecipitation (ChIP) with MLL (**C**) or SETD1A (**D**) and IgG antibodies were performed on HEK-293 cells. Immunoprecipitated DNA was quantified with RT-qPCR and results plotted as percent input enrichment, are shown. (**E-F**) Shown are analyses of ChIP with MLL (**E**) or SETD1A (**F**), and IgG antibodies, performed on IMR-90 tert cells, as described above. (**G-H**) H3K4me2 and IgG chromatin immunoprecipitation was performed on HEK-293 (**G**) and IMR-90 tert (**H**) cells, and the result plotted as percent input enrichment, are shown. Each experiment was performed at least three, or more times. Error bars represent SD. *P ≤ 0.05, **P ≤ 0.005, *** P ≤ 0.0005, ns: not significant P > 0.05 (two-tailed Student’s t-test). α-sat, α-satellite.

To confirm our observations from the IF, we performed chromatin immunoprecipitation (ChIP) using a specific antibody against MLL or SETD1A and checked for their binding on the centromeres in HEK-293 cells (Figure 2C-D). We used their canonical targets i.e. *HOXA9* (& *PAX9*) for MLL and *RAD18* for SETD1A as positive control (Alsulami et al., 2019; Wang et al., 2009). In our ChIP samples, we could detect enrichment of MLL and SETD1A over IgG on both α-satellite regions as well on chromosome 17 (Figure 2 C-D). Independent of their centromere-forming status, we detected MLL and SETD1A on all three HORs in chromosome 17 (Figure 2 C-D). In order to ascertain that the binding of MLL and SETD1A on centromeres is specific, we performed two additional experiments. First, we knocked down MLL (Supplementary Figure S2F) or SETD1A (Supplementary Figure S2H) using shRNAs which enabled us to obtain sufficient cells for our ChIP assay. Consistent with the reduced binding of MLL and SETD1A on *HOXA9* and *RAD18* promoters respectively, we observed that their enrichment was also significantly reduced on the α-satellite loci (Supplementary Figure S2G, I). Second, we performed ChIP in non-transformed IMR90-tert cells, and found that both MLL and SETD1A bound to the α-satellite region and chromosome17 HORs in these cells as well (Figure 2E-F). We simultaneously performed ChIP with CENP-B, a DNA-binding protein that binds to a 17bp-consensus sequence present in α-satellite loci (Morozov., 2017; Masumoto et al., 1989). The CENP-B showed significant enrichment at centromeres but not at other non-centromeric loci (Supplemental Figure S2J), indicating that we are able to amplify and detect centromere enrichment on endogenous chromosomes with our ChIP experiments. In addition to the H3K4 methyltransferase, we observed high levels of H3K4me2 in our ChIP experiments in both HEK-293 (Figure 2G) and IMR90-tert cells (Figure 2G). Altogether, our results show that both the H3K4 depositing enzymes as well as the H3K4 dimethylation marks are present at the centromeres.

### Loss of MLL affects the epigenetic landscape of the centromeres

Previous studies show that the H3K4me2 mark is essential for the transcription as well as the stability of the centromere (Molina et al., 2016; Martins et al., 2016; Bergmann et al., 2011). Here, we have reported the presence of most members of H3K4 HMT family on the centromere (Figure 2, S2E). In order to investigate, if these proteins indeed deposit the H3K4me2 marks on the centromeres, we decided to proceed with the analysis of one of these H3K4 HMT members— MLL— in greater detail. In the studies with HAC, it was observed that the effects of removal of the H3K4me2 mark were apparent after some days, as CENP-A turnover is slow (Molina et al., 2016; Bergmann et al., 2011). Therefore, we decided to generate MLL knock-out cell lines. To achieve this, we performed CRISPR-Cas9 based genome editing on HEK-293 cells to produce MLL knock-out cell lines. Our initial attempts to generate MLL knock-outs in several different cell lines were not successful. Therefore, keeping in mind that MLL is essential for cell viability and growth (Sugeedha et al., 2020; Crump and Milne, 2019), we generated and successfully obtained HEK-293 cell lines using doxycycline-inducible Cas-9 expression vectors (Wang et al., 2014; Figure 3A). MLL levels were drastically reduced in the two independent MLL-knock-out clones shown here (MLL iKO#11 and iKO#20). In the case of chromatin-binding proteins, often cellular levels show reduction but not the chromatin-bound fraction. We, therefore, confirmed that MLL chromatin binding was indeed reduced in our inducible knock-out cell lines by ChIP assays (Figure 3B, S3A), both on centromere and *HOXA9* promoter. After successfully replicating these ChIP experiments in the iKO cell lines several times, we interrogated the effect of loss of MLL on H3K4 dimethylation levels. As expected, the H3K4me2 levels were dramatically reduced in both MLL iKO cell lines (Figure 3C, S3B), indicating that MLL was indeed one of the writers of H3K4me2 at the centromeres. Consistent with the observations on HAC, reduction in the H3K4me2 mark was accompanied by a reduction in H3K9 acetylation as well as an increase in H3K9me3 (Figure 3D-E, S3C-D). Surprisingly, despite reduced transcription upon loss of MLL, we did not observe a decrease in the level of the dimethylation at H3K36, rather it showed an overall increase (Figure 3F, S3E). These observations are in contrast with the results obtained in HAC, where the H3K36me2 mark was reduced upon lysine-specific demethylase1/2 (LSD1/2) targeting (Molina et al., 2016; Bergmann et al., 2011). Notably, the H3K36me2 mark was only increased at the centromere MLL iKO cells but not at the canonical locus —*PAX9* promoter (Figure 3F, S3E). All in all, these observations indicate that MLL regulates the local epigenetic landscape of centrochromatin by regulating the levels of H3K4 dimethylation marks.

**Figure 3:**
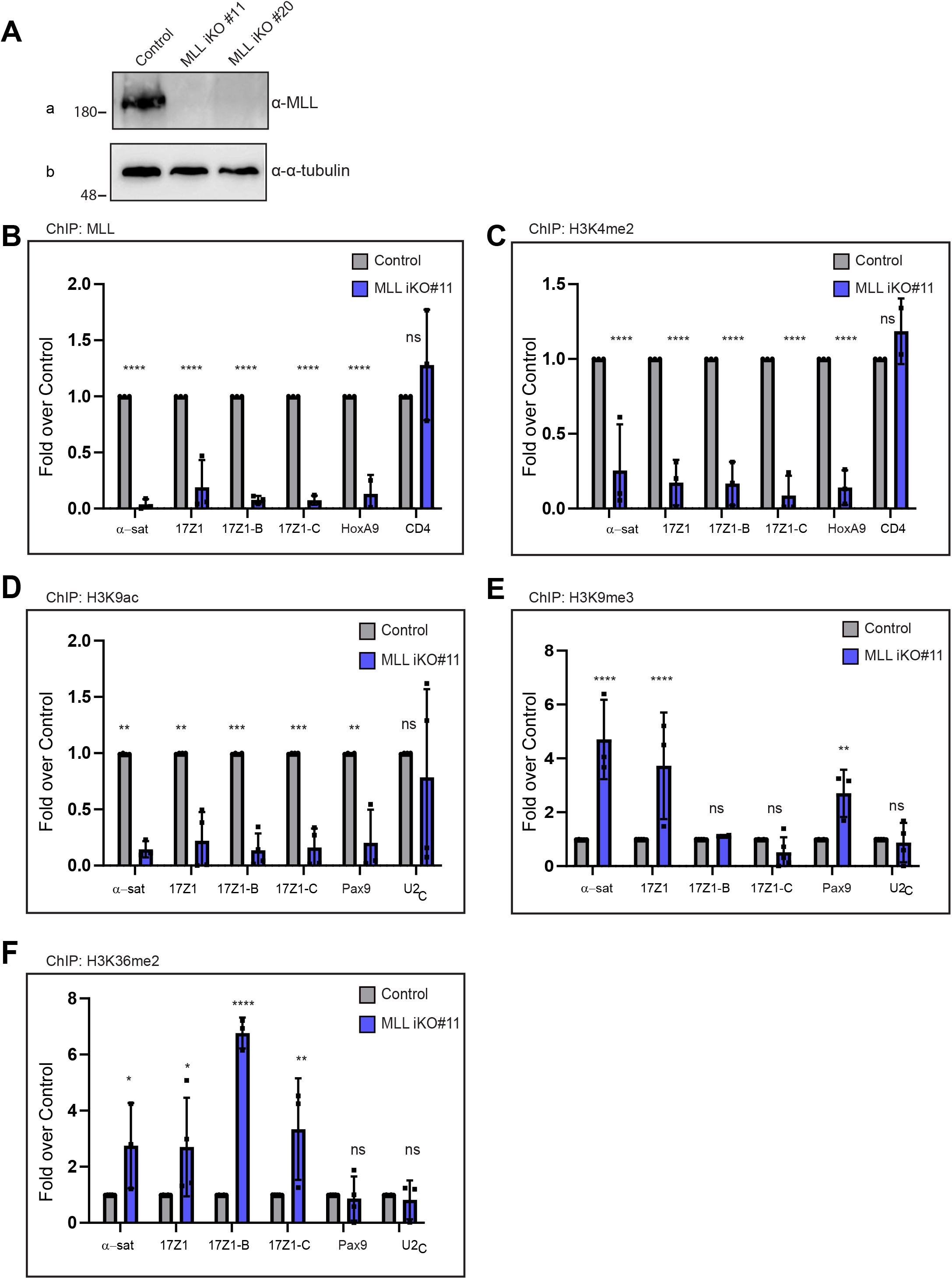
Loss of MLL affects the epigenetic landscape of the centromeres. (**A**) Immunoblot shows MLL protein levels in CRISPR-Cas9 generated inducible *MLL* knockouts (iKO) cells. Two independent clonal cell lines (#11 and #20) were used here. Blots were probed with α-MLL_C_ and α-tubulin as shown (*Please note that the same sample was loaded on a different SDS-PAGE gel to evaluate tubulin*). Molecular weight markers (in kDa) are shown on the left. (**B**-**F**) ChIP analyses using following antibodies: MLL (**B**), H3K4me2 (**C**), H3K9ac (**D**), H3K9me3 (**E**), and H3K36me2 (**F**) in *MLL* iKO cells (#11) are shown. Data were normalized against the ChIP values obtained in parental (or Cas9-expressing) cells, which are used as Control. Each experiment was performed at least three, or more times. Error bars represent SD. *P ≤ 0.05, **P ≤ 0.005, *** P ≤ 0.0005, ****P ≤ 0.0001, ns: not significant P > 0.05 (Two-way ANOVA with Šídák multiple comparison test). α-sat, α-satellite.

### Disparate impact of MLL and SETD1A on centromeric R-loops

Recently, several studies have reported the presence of R-loops on the centromere, which have been shown to affect centromeric stability (Racca et al., 2021; Giunta et al., 2021; Mishra et al., 2021; Kabeche et al., 2018). As MLL is involved in the transcription of cenRNA, we asked if loss of MLL would influence the status of R-loops on the centromere. To this end, we first used the S9.6 antibody to detect global nuclear R-loops by IF. The R-loop signal intensity was quantified by measuring the mean signal intensity in the nucleus in Control Vs. Test cells. When quantified, to our surprise, we found that loss of MLL resulted in a higher R-loop signal compared to control U-2OS cells (Figure 4A, S4A). Given MLL’s role in cenRNA transcription, this result was unexpected. Further, a recent study reported the loss of R-loops upon SETD1A siRNA treatment (Yilmaz et al., 2021). Therefore, we quantified R-loop levels in SETD1A siRNA-treated cells. In contrast to MLL and consistent with the previous report, loss of SETD1A resulted in lower R-loop formation compared to Control cells (Figure 4A). RNA Pol II inhibitors are known to reduce R-loop formation due to inhibition of transcription (Racca et al., 2021). We found that treatment with Triptolide diminished R-loop formation in MLL siRNA treated cells, when compared with Triptolide treated Control siRNA samples (Figure 4A, compare sample 4 with 5) or MLL siRNA non-Triptolide treated samples (Figure 4A, compare sample 2 with 5). However, we observed no significant change between SETD1A-siRNA non-Triptolide Vs Triptolide treated samples (Figure 4A, compare sample 3 with 6). These results indicate that the primary reason for the increase or decrease of R-loops upon loss of our HMTs is transcription. To validate our findings of R-loops in IF, we performed DNA:RNA hybrid immunoprecipitation (DRIP) assay in MLL or SETD1A siRNA-treated cells. The RNase H treated genomic DNA before the DRIP assay worked as a control to ensure a specific signal (Sridhara et al., 2017). An R-loop-prone locus – *RPL13A* (intron 7 and exon 8; Sanz et al 2019), acted as a positive control, while R-loop-free loci – *SNRPN* and *EGR1* (Sridhara et al 2017, Sanz et al 2019), and MLL negative locus—U2_C_ region (Zargar et al., 2018) acted as negative controls (Supplementary Figure S4B). Consistent with our IF data, loss of MLL increased R-loop signal by several folds on the centromere (Figure 4B) whereas SETD1A siRNA treatment resulted in a reduction of centromeric R-loop formation (Figure 4C). The DRIP signal was significantly reduced by pre-treatment with RNase H in all cases (Control, MLL, and SETD1A siRNA) indicating that we can reliably detect specific R-loops in our experiments. To sum up, our results show that MLL and SETD1A behave differently in regulating R-loop formation on the centromere and their ability to promote/reduce R-loop accumulation is associated with transcription.

**Figure 4:**
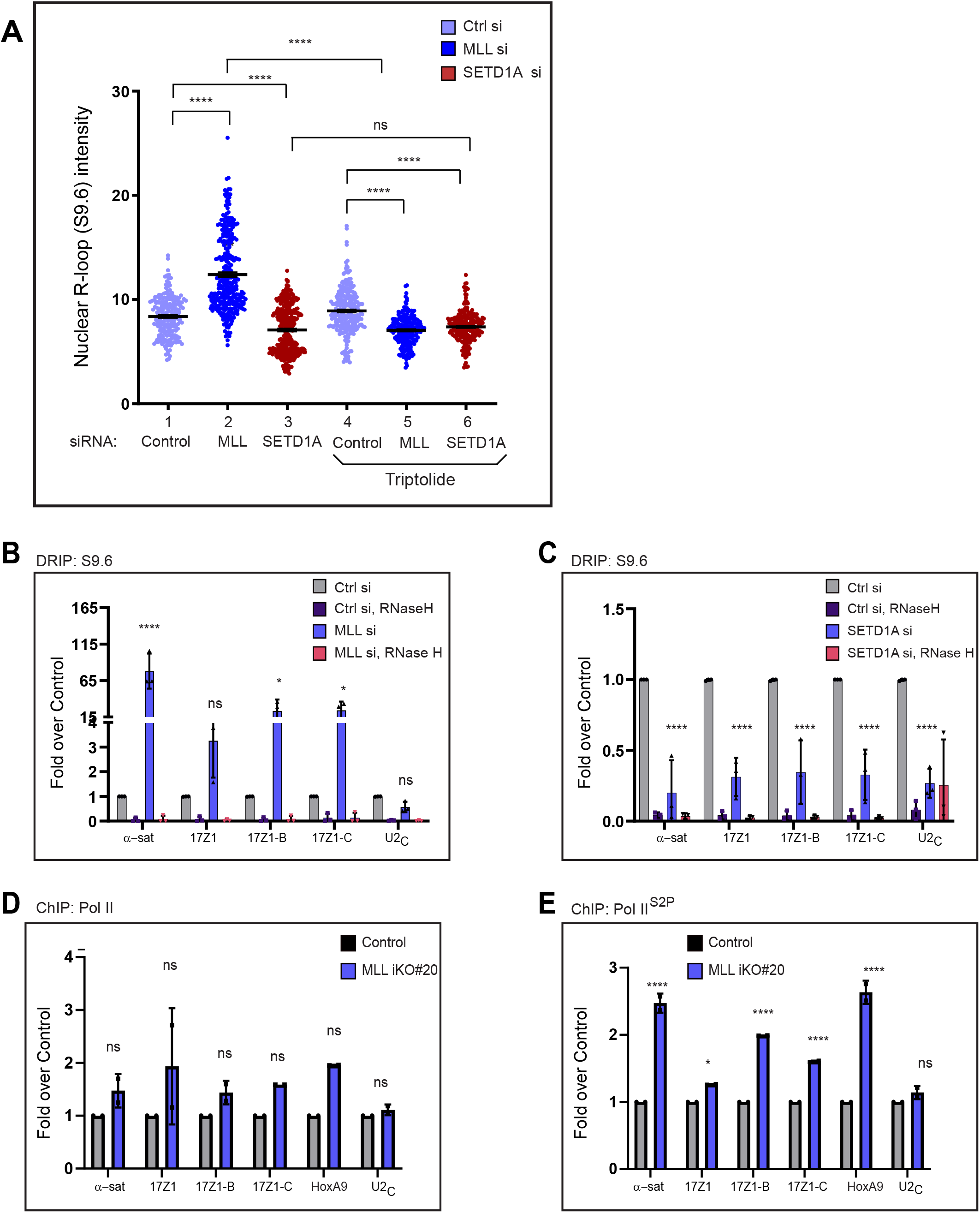
Disparate impact of MLL and SETD1A on centromeric R-loops. (**A**) Quantification of nuclear R-loops in U-2OS cells stained by S9.6 antibody, 48 hrs after Control, MLL, or SETD1A siRNA treatment, is shown. For transcription inhibition, cells were treated with 20 µM triptolide or DMSO (control) for 4 hrs. The intensity of the whole nuclear R-loop staining is plotted. A total of 100 cells from three independent experiments was scored. See Figure S4A for representative images. Error bars represent SEM. ****P ≤.0001, ns: not significant P>0.05 (Mann-Whitney two-tailed unpaired test). (**B**-**C**) DNA:RNA immunoprecipitations (DRIP) after MLL (**B**) and SETD1A (**C**) RNAi in HEK-293 cells, with respective RNase H controls, is shown. Data were normalized against the DRIP values obtained in Control siRNA-treated cells (Ctrl si). Note that the *U2_C_* control (in B) is the same data from Figure S4B but normalized against DRIP values obtained in Control cells. Each experiment was performed at least three, or more times. Error bars represent SD. *P ≤ 0.05, ****P ≤ 0.0001, ns: not significant P > 0.05 (Two-way ANOVA with Šídák multiple comparison test). (**D**-**E**) ChIP-analysis of RNA Pol II (**D**) and RNA Pol II^S2P^ (**E**) in *MLL* iKO#20 cells are shown. Data were normalized against the ChIP values obtained in parental (or Cas9-expressing) cells, which are used as Control. Data from two independent ChIP experiments are plotted. Error bars represent SD. *P ≤ 0.05, ****P ≤ .0001, ns: not significant P > 0.05 (Two-way ANOVA with Šídák multiple comparison test). Ctrl, control; si, siRNA.

### Loss of MLL perturbs RNA Pol II distribution at the human centromeres

Centromeres are known to be transcribed by RNA Pol II and R-loops are a by-product of transcription (Corless et al., 2020; Perea-Resa and Blower, 2018; Aguilera and García-Muse, 2012). In order to understand, why R-loops are accumulating upon loss of MLL but not SETD1A, we stained for RNA Pol II on the centromere. We looked at total RNA Pol II (Supplementary Figure S4C) and elongating RNA Pol II as scored by RNA Pol II phosphorylated on CTD serine 2 (RNA Pol II^S2P^) (Supplementary Figure S4D), via IF staining on centromere after treating the cells with MLL or SETD1A siRNAs. We observed that there was little effect of MLL siRNA on the presence of both forms of RNA Pol II. In contrast to these observations and consistent with the model generated by HAC, where H3K4me2 facilitates RNA Pol II-mediated transcription, loss of SETD1A exhibited dispersed foci for both forms of RNA Pol II and, even displayed reduced intensity for RNA Pol II^S2P^ (Supplementary Figure S4C-D). We quantified the intensities of total RNA Pol II and observed an increase in intensity in both MLL and SETD1A siRNA-treated samples (Supplementary Figure S4E). This was probably due to the high background staining present around CENP-A in SETD1A siRNA-treated samples, even though clear intense foci were absent for corresponding RNA Pol II images (Supplementary Figure S4C). In contrast, RNA Pol II^S2P^ showed increased intensity upon MLL siRNA treatment and decreased intensity upon SETD1A siRNA treatment as reflected in the images (Supplementary Figure S4D, F). Previous reports indicate that loss of MLL may result in abnormal distribution of Pol II at a subset of genomic loci (Miyamoto et al., 2020; Wang et al., 2009; Milne et al., 2005). In order to understand what is happening at the centromeres, we performed ChIP with total RNA Pol II and RNA Pol II^S2P^ in MLL iKO cells. Our analysis revealed that the levels of total RNA Pol II did not show any significant change in Control Vs. MLL iKO samples (Figure 4D, S4G). However, we did find increased levels of RNA Pol II^S2P^ accumulated on the centromere upon loss of MLL (Figure 4E, S4H). Taken together, our results show that although MLL and SETD1A deposit H3K4me2 at the centromeres (Figure 3C; Yilmaz et al., 2021), they may differentially regulate RNA Pol II distribution, at least at centromeres. It is presumably this difference that is responsible for the increase in R-loops upon loss of MLL.

### MLL and SETD1A affect kinetochore maintenance and the recruitment of CENP-B and CENP-C to the centromere

Centromere transcription has been implicated in kinetochore maintenance as cenRNA is required for accurate localization of many centromere-associated proteins including CENP-C (McNulty et al., 2017; Rosic et al., 2014). Recently, CENP-B was also shown to be bound by transcripts from inactive arrays (McNulty et al., 2017; Morozov et al., 2017). As both MLL and SETD1A affect the transcription of cenRNA from active as well as inactive arrays, we analysed the effect of MLL or SETD1A knockdown on the kinetochore maintenance. When we checked the localization of CENP-C and CENP-B on the centromere by IF, we observed that the centromeric levels of both CENP-C and CENP-B proteins were decreased upon MLL or SETD1A siRNA treatment (Figure 5A-B). We further analysed these protein levels on the centromere in cell lines expressing different domain deletion of MLL or SETD1A upon siRNA treatment, and as shown, we observed a consistent decrease in both CENP-C (Figure 5A, C-D) and CENP-B (Figure 5B, E-F) on the centromere. We also made the following observations during these experiments: (i) we observed an increase in the levels of CENP-C but not CENP-B in cell lines expressing full-length MLL or SET1A protein (Figure 5C-D, E-F). This can be partly explained due to an increase in cenRNA transcript observed upon expression of SETD1A full-length protein (Figure 1E), indicating that centromeric transcripts indeed play a role in recruiting/stabilizing CENP-C to the centromeres; (ii) the U-2OS cell lines expressing the HMT mutants behaved differently than U-2OS parent cells in terms of staining for centromere proteins. We had difficulty staining for all centromere proteins described in Figure 5 and 6 in our mutant cell lines. This can indicate that mutants of MLL or SETD1A used here, exert a dominant-negative effect on the localization and /or staining of centromere proteins; and (iii) cellular levels of all proteins analyzed here (in Figure 5 and 6) remain largely unchanged upon loss of HMT, except CENP-B, which exhibited decreased levels (Supplementary Figure S5 A-D). Further analysis using previously published data sets (Miyamoto et al., 2020; Wang et al., 2009) indicated that CENP-B may be a direct transcriptional target of MLL. Our results indicate that MLL family regulates the kinetochore maintenance at several levels—in *cis* by modulating cenRNAs at centromeres (and therefore impacting the recruitment of centromere proteins (like CENP-C) or in *trans* by regulating transcription of centromeric/kinetochore gene (like *CENP-B*). All in all, loss of MLL and SETD1A have an adverse effect on the maintenance of kinetochores.

**Figure 5:**
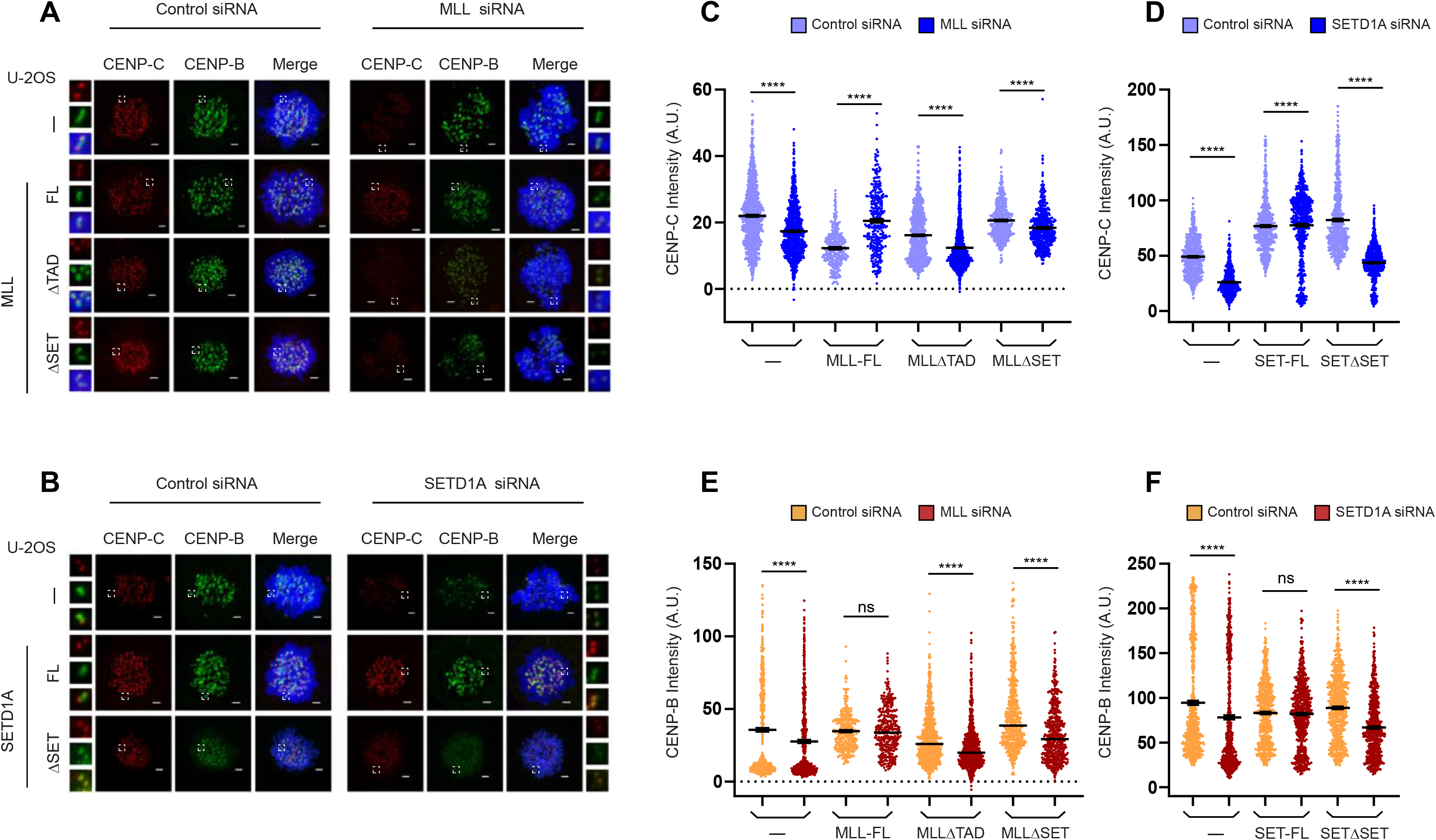
MLL and SETD1A affect kinetochore maintenance. (**A**) Representative IF images show mitotic CENP-C (red) and CENP-B (green) staining in parent U-2OS cells (—) or U-2OS cells expressing full-length MLL (MLL-FL), TAD deleted MLL (MLLΔTAD) and SET domain deleted MLL (MLLΔSET) following treatment with either Control or MLL siRNA. (**B**) Representative IF images show mitotic CENP-C (red) and CENP-B (green) staining following treatment with Control or SETD1A siRNA in parent U-2OS cells (—) or U-2OS cells expressing full-length SETD1A (SET-FL), and SETD1AΔSET (SETΔSET; here N1646A mutant was used). (**A**-**B**) DNA was stained with DAPI (blue). The area in the white square is magnified and shown on the side of each image. Scale bar, 2μm. (**C**-**E**) The graph represents the quantification of CENP-C (**C**) and CENP-B (**E**) intensity in MLL depleted cells shown in **A**. (**D**-**F**) Quantification of CENP-C and CENP-B intensity following treatment with SETD1A siRNA as shown in **B**. (**C**-**F**) Each data point represents a single centromere. The error bar represents SEM. n ≥300 centromeres (n=2 experiments). For quantification, Z-stack images were merged and individual CENP-C and CENP-B signal intensity were measured using ZEN software. ****P ≤ 0.0001, ns: not significant P > 0.05 (Mann-Whitney two-tailed unpaired test).

**Figure 6:**
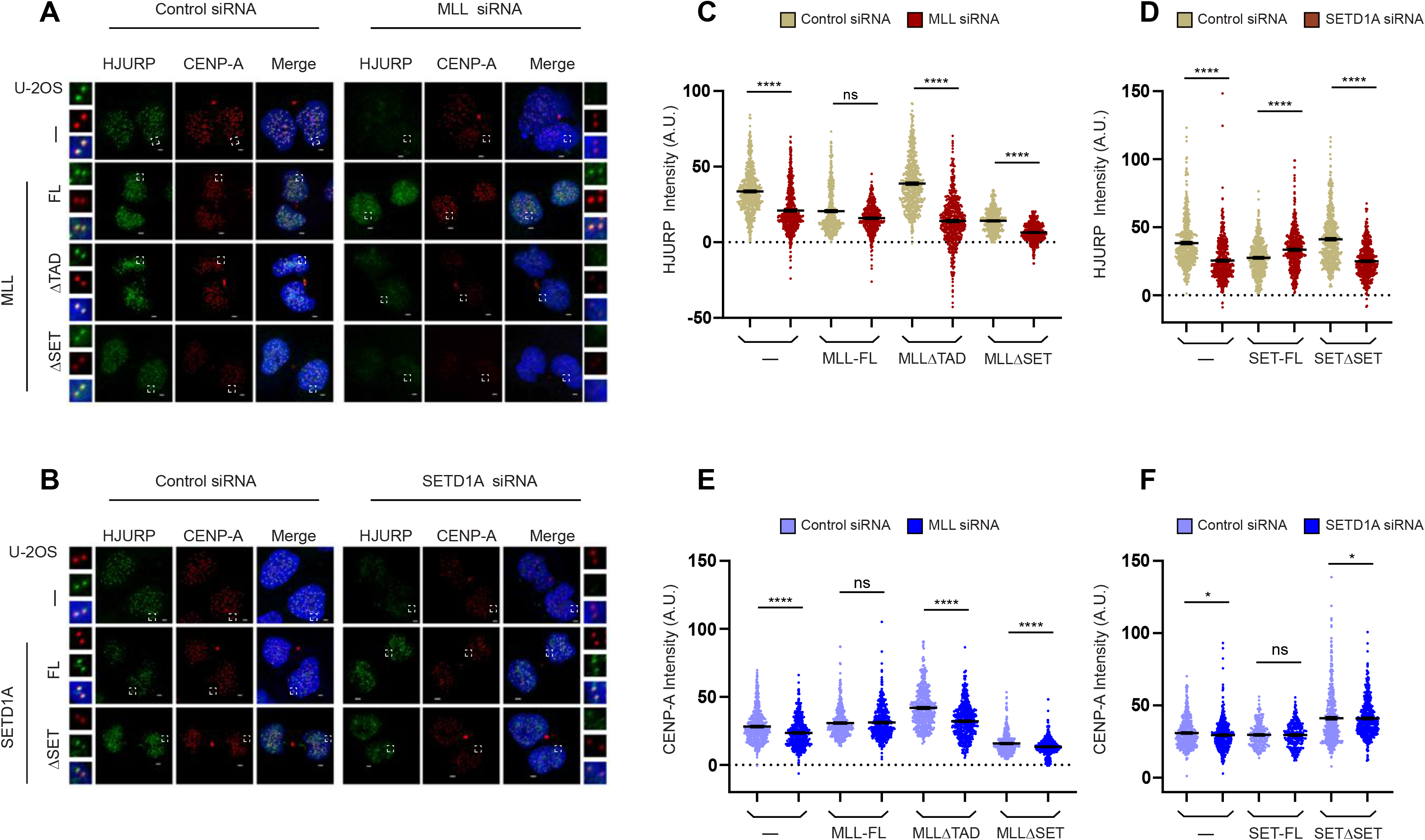
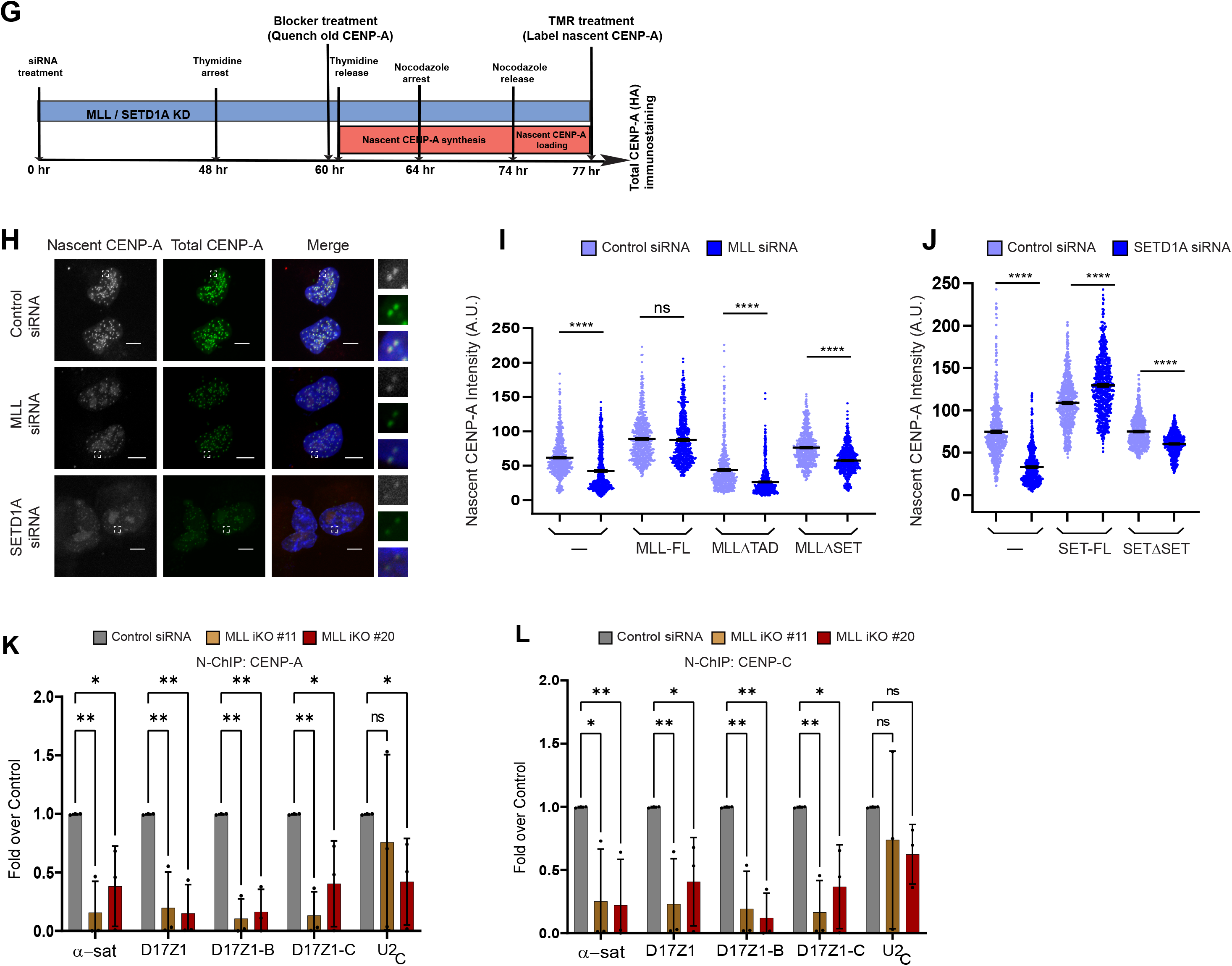
MLL and SETD1A facilitate recruitment of nascent CENPA at centromeres. (**A**) Parent U-2OS cells (—) or U-2OS cells expressing recombinant full-length MLL (MLL-FL), TAD deleted MLL (MLL-ΔTAD), and SET domain deleted MLL (MLL-ΔSET), were treated with either Control or MLL siRNA. The cells, synchronized for the early G1 phase, were stained with α-CENP-A (red) and α-HJURP (green) antibody as shown. (**B**) Parent U-2OS cells (—) or U-2OS cells stably expressing recombinant full-length SETD1A (SET-FL), and SET domain deleted SETD1A (SETΔSET) were treated with either Control or SETD1A siRNA. The representative IF images show the loading of HJURP and CENP-A at the centromere in early G1 phase cells. (**A**-**B**) DNA was stained with DAPI (blue). The area in the white square is magnified and shown on the side of each image. Scale bar, 2μm. (**C**-**E**) Quantification of HJURP and CENP-A fluorescence intensity following depletion of MLL (**C**, **E**) and SETD1A (**D**, **F**) respectively. Each data point represents a single centromere. The error bar represents SEM. ≥250 centromeres quantified from 10 early G1 cell pairs, (n=2 experiments). ****P ≤ 0.0001, *P ≤ 0.05, ns: not significant p > 0.05 (Mann-Whitney two-tailed unpaired test). (**G**) Schematic of cell synchronization and TMR-based labelling strategy to detect nascent CENP-A upon MLL/SED1A knockdown is shown. See methods for more details. (**H**) IF images showing the effect of Control/MLL/SETD1A siRNA treatment on nascent CENP-A loading at the centromere in U-2OS cells are shown. Cells stably expressing the CENP-A-SNAP-HA construct were used for siRNA treatment. Nascent CENP-A was labelled with TMR (grey) while total CENP-A was detected by IF using α-HA antibody (green). The area in the white square is magnified and shown on the left for each image. Scale bar, 5μm. (**I**) Quantification of centromeric fluorescence intensity of nascent CENP-A-SNAP in parent U-2OS cells (—) or cell line stably expressing siRNA resistant MLL full length (FL) or MLLΔTAD or MLLΔSET upon MLL siRNA. (**J**) Quantification of centromeric fluorescence intensity of nascent CENP-A-SNAP in parent U-2OS cells (—) or cell line stably expressing siRNA resistant SETD1A full length (FL) or SETD1AΔSET (here N1646A mutant was used) upon SETD1A siRNA is shown. (**I**-**J**) Each data point represents a single centromere. Maximum Intensity projection (MIP) images were used in all quantification analyses using ZEN software. n ≥ 300 quantified from 10 early G1 cell pairs, (n=2 experiments). ****P ≤ 0.0001, ns: not significant p > 0.05; (Mann-Whitney two-tailed unpaired test). (**K**-**L**) Shown are native ChIP-analysis of CENP-A (**K**) and CENP-C (**L**) in *MLL* iKO cells (#11 and #20). Data were normalized against the ChIP values obtained in parental (or Cas9-expressing) cells, which are used as Control. Data from three or more independent ChIP experiments are plotted. Error bars represent SD. *P ≤ 0.05, **P ≤ 0.005, ns: not significant P > 0.05 (Two-way ANOVA with Šídák multiple comparison test). A.U., arbitrary units.

### MLL and SETD1A facilitate recruitment of nascent CENPA at centromeres

Both CENP-C and CENP-B participate at different levels to stabilize CENP-A nucleosomes (Carty et al., 2021; Otake et al., 2020; Falk et al., 2015; Fachinetti et al., 2015). Further, the recruitment of CENP-A chaperone HJURP is dependent on H3K4me2-facilitated transcription (Bergmann et al., 2011). Hence, we sought to determine if the recruitment of HJURP on the centromere, and as a consequence that of CENP-A, is affected upon loss of our H3K4 HMTs. We depleted MLL or SETD1A in our parent U-2OS cell lines and co-stained the cells for endogenous HJURP and CENP-A in early G1 cells (Figure 6 A-B). Consistent with results observed with CENP-C, HJURP and CENP-A recruitment was diminished in parent U-2OS cells as well as cells expressing MLLΔTAD and MLLΔSET treated with MLL siRNA, but not full-length MLL (Figure 6A,C,E). Similarly, upon treatment with SETD1A siRNA, parent U-2OS cells and SETD1AΔSET cell lines showed significant loss of HJURP and CENP-A on the centromere, while expression of full-length SETD1A was able to restore their levels (Figure 6B,D,F). In order to confirm that the loss of CENP-A observed here is not due to reduced cell proliferation upon loss of MLL or SETD1A (Ali et al., 2014; Salz et al., 2014; Liu et al., 2010) we made use of the pulse-chase labeling approach by using SNAP tagged CENPA (Jansen et al., 2007) which can reveal if incorporation of newly synthesized CENP-A is affected or not. We have shown that treatment with MLL or SETD1A siRNAs resulted in a 50% reduction in cenRNA after 72 hrs (Figure 1A). At this time at least 50% of cells are still division-competent (Ali et al., 2014). Therefore, 60 hrs after siRNA treatment, the old CENP-A was blocked and newly synthesized CENP-A loading was detected by TMR-staining in cells synchronized in early G1 phase (Figure 6G). As shown, we observed about 50-60% reduction in nascent CENP-A loading upon perturbing MLL or SETD1A levels (Figure 6 H-J). Interestingly, this decrease was restricted to 15-25% upon compromising the HMT domains of MLL or SETD1A (Figure 6 I-J). On the other hand TAD deletion in MLL caused about 42% decrease in nascent CENP-A levels, much nearer to the levels observed in parent U-2OS cells (Figure 6I). Our findings suggest that inhibition of potent transcription, like by the TAD, shows a more immediate effect on nascent CENP-A loading than depletion of H3K4me2, whose effects may be manifested over a longer period of time. Remarkably, we also observed a decrease in the exogenously expressed total CENP-A levels on the centromere when their protein levels were unchanged (Supplementary Figure S5E-H). Our results indicate that MLL or SETD1A depletion was more deleterious than RNAi-mediated depletion of cenRNA (McNulty et al., 2017; Quénet and Dalal, 2014) or LSD1/2 -mediated removal of H3K4me2 (Molina et al., 2016; Bergmann et al., 2011). Encouraged by our observations here, we performed native ChIP using CENP-A and CENP-C antibodies in MLL iKO cells. Consistent with our observations in IF, the level of CENP-A and CENP-C showed a reduction on α-satellite loci in cells devoid of MLL (Figure 6 K-L, S5 A,D). Altogether, our results indicate that MLL and SETD1A modulate the epigenetic state of the centromere by their histone methyltransferase/transcription activity to regulate the cellular machinery involved in CENP-A deposition.

## Discussion

The discovery of active histone mark – H3K4me2 – within the centrochromatin and the elegant demonstration of its crucial role in kinetochore maintenance using targeted-engineering of HAC, has firmly established the importance of this mark in kinetochore function. However, experimental evidence identifying which of the many histone lysine methylation enzymes deposits this mark on the centromere, was lacking. Despite differences in size, structure, interacting partners, and catalytic potential, the various members have been grouped under the MLL (KMT2) family banner by virtue of their SET domain. Even though each of these proteins is uniquely required during development, redundant roles of these enzymes are well known (Sugeedha et al., 2020; Piunti and Shilatifard., 2016). In this study, we have shown that the majority of MLL family members associate with the centromeres and regulate centromeric transcription. Our study here highlights that not only the H3K4me2 mark but the enzyme depositing it, also contributes to centromere stability.

### MLL regulates kinetochore function in multiple ways

Our results suggest that MLL affects the kinetochore assembly and maintenance in multiple ways. Not only is the presence of the SET domain in MLL required at the centromere to deposit the H3K4me2 mark, but it also co-activates centromeric transcription by its TA domain. Both these activities are pertinent in recruiting key proteins like CENP-C and HJURP to the centromere and therefore loading of CENP-A at the centromere. In fact, our results indicate that loss of rapid transcription (by deletion of TAD) exhibits pronounced effect on *de novo* CENP-A incorporation in our assay, even though the decrease in cenRNA transcripts by loss of both TAD and SET-domain deletion is comparable (Figure 1, 6I-J). MLL also regulates the cellular levels of proteins like CENP-B, which are important for centromere function. Overall, loss of CENP-A is tolerated by the cell with chromosome segregation defects appearing after multiple rounds of cell division (McNulty et al., 2017; Molina et al., 2016; Bergmann et al., 2011). We have previously reported modest chromosome mis-alignment upon loss of SET or TA domain (Ali et al., 2017). Indeed, mutation of the WDR5-interacting motif in MLL turned out to be the major cause of chromosome misalignment in our assays after 72 hr. of RNAi. However, we also believe that MLL participates in multiple pathways to regulate chromosome segregation. This statement is prompted by our earlier observation that about 90% of all MLL-depleted cells showed segregation defects but only 25% of these cells showed elongated phenotype which can be attributed to loss of Kinesin-like protein 2A (KIF2A) function (Ali et al., 2017). Furthermore, we also observe problems in centrosome function upon loss of MLL, which can contribute to defects in spindle formation and therefore, chromosome segregation (Chodisetty and Tyagi, unpublished data). Studies from the HAC model have revealed that kinetochores can function for several rounds of cell division without displaying any prominent defects even after the loss of H3K4me2, and with only 50% CENP-A on the centromere (Molina et al., 2016; Bergmann et al., 2011). Our assays for nascent CENP-A deposition, though effective over multiple arrays, manifest loss of 50% CENP-A after 72 hr. of RNAi. Remarkably, this loss is only 25% in MLLΔSET mutant indicating that these cells will take much more time to exhibit failure of kinetochore activity than can be addressed in our current assay. This could explain why we did not detect segregation defect as a primary phenotype of SET-domain deleted mutants in our earlier reports. Alternatively, the redundant functional activity of family members and/or alternate pathways could circumvent the loss of MLL. For example, yeast Set1A has been shown to regulate spindle assembly checkpoint through its interaction with mitotic arrest deficient 2 (Mad2) and regulate Ipl1-Aurora kinase by methylating outer-kinetochore protein Dam1 (Schibler et al., 2016; Zhang et al., 2005); both processes ensure the proper segregation of chromosomes during mitosis. All in all, the regulatory relationship of the Set1A/MLL family members on the centromere is undeniable.

### Epigenetic landscape of human endogenous centromere: differences from the HAC model

Our quest on understanding mitotic roles of MLL highlights the fact that loss of these proteins has a pleiotropic effect on cellular processes making it hard to attribute one defect to one process. For such proteins, the use of HAC is an effective tool in teasing out mechanistic details of one process at a time on the centromere. On the flip side, till the cross-talk of multiple pathways is not appreciated, the cumulative impact of a histone modifier cannot be correctly gauged. For instance, in line with observations in HAC, we observed lower levels of active transcription marks (H3K4me2 and H3K9ac) and an increase in the inactive transcription mark (H3K9me3) on the centrochromatin in MLL iKOs. In contrast to the reports in HAC (Molina., 2016; Bergmann et al., 2011), we observed an increase in the transcription elongation mark H3K36me2 in MLL iKO cells at the centromere. Without doubt, H3K36me2 mark is primarily associated with active transcription (Li et al., 2019; Krogan et al., 2003; Hui et al., 2003). In agreement, besides above-mentioned reports, another study observed that loss of KDM2A, the H3K36me2 demethylase, is associated with higher α-satellite transcription (Frescas et al., 2008). However, different from reports in HAC, we also observed accumulation of RNA Pol II ^S2P^ (and R-loops) in MLL iKOs (also see below). An increase in aberrant R-loops can lead to DNA double-strand breaks at several genomic locales and challenge centromere integrity (Castillo-Guzman et al., 2021; Giunta et al., 2021; Racca et al., 2021). Interestingly, H3K36me2 increases in cells with exaggerated DNA double-strand breaks (Huan et al., 2018; Cao et al., 2016; Fnu et al 2010), a phenomenon which has been reported on centromeres (Guinta et al., 2021;Yilmaz et al., 2021). Hence, our data can be better explained considering the nature of H3K36me2 both as a transcription elongation mark and a DNA break repair factor, a possibility that needs further testing.

### MLLs regulate co-transcriptional R-loops at the centromere

Previous studies report a positive correlation between transcription, H3K4me2 mark and R-loop formation (Sanz et. al., 2016; Chen et al., 2015) and R-loops at centromere are no different (Yilmaz et al., 2021). Our data with SETD1A knock-down confers with this model and showed a decrease in R-loop formation. In contrast, loss of MLL, despite showing decreased levels of H3K4me2 on the centromere, gave rise to an increase in R-loop formation. Even though, this MLL-RNAi based R-loop formation was transcription-dependent, we observed a concomitant increase in RNA Pol II ^S2P^ at the centromere depicting an aberrantly ‘paused’ RNA Pol II in MLL iKO cells. Indeed, a stalled RNA pol II is associated with R-loop formation (Sridhara et al., 2017). Due to our inability to generate SETD1A knock-out cells, we are unable to interrogate the status of RNA Pol II^S2P^ in the absence of SETD1A on the centromere. However, studies from yeast, *Saccharomyces cerevisiae*, show that deletion of the only H3K4 methyltransferase (*Set1A*) in the cell has little effect on the recruitment of RNA Pol II (Krogan et al., 2003; Hui et al., 2003). This process may be conserved in higher organisms as SET domain deleted mutant of SETD1A showed no compelling changes in RNA Pol II occupancy in mammalian cells (Sze et al., 2017 ; Lee and Skalnik, 2008). In contrast, loss of function of MLL results in varied defects in RNA Pol II distribution (Miyamoto et al., 2020; Wang et al., 2009; Milne et al., 2005). While reduced RNA Pol II occupancy has been reported at some loci, an increase in RNA Pol II^S2P^ and serine 5-phosphorylated RNA Pol II (RNA Pol II^S5P^) forms have been reported at others (Wang et al., 2009; Milne et al., 2005). A recent study shows an increase of RNA Pol II levels at transcription termination sites in MLL KO cells (Miyamoto et al., 2020) indicating that abnormal distribution of Pol II can ensue following a loss of MLL. Here we report a novel role of MLL in R-loop formation on the centromere. Our study also highlights the fact that despite being such well-studied co-activators of RNA Pol II, how different members of the MLL family regulate RNA Pol II is still not clear.

R-loops reported at centromeres have been proposed to be beneficial (Yilmaz et al., 2021; Racca et al., 2021; Kabeche et al., 2018) as well as detrimental (Giunta et al., 2021; Mishra et al., 2021) to centromere integrity. Recent reports suggest that CENP-A and Aurora B Kinase can prevent the formation of opportunistic R-loops at centromeres in a cell-stage-specific manner (Guinta et al., 2021; Moran et al., 2021). Furthermore, we found that loss of both MLL and SETD1A severely impacts CENP-A loading at the centromeres. While the loss of MLL and SETD1A can trigger replication stress and DNA damage (Hoshii et al., 2018; Liu et al., 2010) which are associated with deleterious R-loops (Castillo-Guzman et al., 2021), how R-loop imbalance at centromeres challenges DNA integrity is still an open question. Altogether, our work raises an interesting hypothesis that MLL and SETD1A may regulate different classes of R-loops (Castillo-Guzman et al., 2021), and thus impact centromere integrity.

### Implications of centromeric transcription regulation in MLL-rearranged leukemias

Growing body of evidence suggest perturbed centromeric and pericentromeric transcription in pathological conditions like cancer (Corless et al., 2020; Ting et al., 2011; Eymery et al., 2009). For example, in lung cancer and squamous cell carcinoma, dysregulation of centromeric transcription was observed accompanied by a global loss of repressive epigenetic marks (Eymery et al., 2009). Similarly, loss of chromatin regulatory proteins has been reported to induce centromeric transcription as a cause or consequence of oncogenesis (Huang and Zhu 2018; Frescas et al., 2008). Furthermore, a decrease in centromeric transcription is lethal to the cell as the absence of tumor suppressor Pbx-regulating protein-1 (Prep1) leads to an increase in repressive marks, resulting in centromere instability (Iotti et al., 2011). Additionally, kinetochore proteins like CENP-K and KNL-1 have been reported as fusion partners of MLL in leukemia (Marschalek, 2016). Intriguingly, long non-coding (lnc) RNAs seem to play a crucial role in MLL-mediated gene regulation (Statello, 2021; Schwarzer et al., 2017). For example, HOTTIP is a well-studied lnc RNAs that interacts with MLL-WDR5 and regulates transcription of *HOXA*-gene cluster through looping of chromatin in normal cells, failure of which could trigger leukemogenesis in mice. Another study reports that lnc RNA UMLILO interacts with MLL-WDR5 and imparts trained immunity in mice (Statello, 2021). Here we found that MLL is regulating the expression of cenRNAs, however, further studies understanding the role of MLL-fusions in centromeric transcriptions still need to be undertaken.

## Methods

### Cell culture and stable cell line generation

U-2OS (human osteosarcoma), HEK-293 (human embryonic kidney), and IMR-90 tert (human lung fibroblast) cells were grown as monolayers in DMEM, supplemented with 10% (v/v) fetal bovine serum (FBS), 1% (v/v) GlutaMAX, and 100 U/mL penicillin-streptomycin. The cells were maintained at 37° C in a humidified atmosphere with 5% CO_2_. All cell lines were authenticated by Lifecode Technologies Private Limited (India).

### Cloning and Site-directed Mutagenesis

U-2OS cells expressing MLL mutants have been described before (Ali et al., 2014) except MLLΔTAD which was generated in full-length MLL here by deletion of aa 2847–2855 using site-directed mutagenesis. Full-length SETD1A cDNA, gift from Robert Roeder (Tang et al., 2013), was cloned in Xho1 linearised pcDNA5/FRT-SFB vector (Zargar et al., 2018). We then generated siRNA resistant full-length SETD1A by introducing seven silent mutations in the full-length construct using site-directed mutagenesis (see Supplemental Figure S1G). The siRNA resistant full-length SFB-tagged SETD1A construct was further used to generate SETD1AΔSET (Δaa1407-1707) and SETD1A N1646A plasmids using site-directed mutagenesis*. MLL* sgRNA were cloned into a lentiGuide-Puro vector, gift from Feng Zhang (Sanjana et al., 2014) in the BsmB1 site. CENPA-SNAP-3xHA ORF, gift from Lars Jansen (Jansen et al., 2007) was cloned into EcoR I and Kpn I linearized pcDNA 3.1-Puro vector or Hind III and Xho I linearized pcDNA FRT vector (Thermo Fischer Scientific).

### Generation of Stable Cell Lines

Cell lines were generated by transfecting the SETD1A constructs using polyethylenimine (PEI; Polysciences Inc.) as described earlier (Zargar et al., 2018). Plasmid transfected cells were selected in the media supplemented with 200μg/ml Hygromycin B (Thermo Fischer Scientific). Cells cultured from individual colonies were used for further experiments. To generate inducible knockouts for MLL, HEK-293 cells were transduced with Doxycycline-inducible Cas9 expression vector pCW-Cas9, gift from Eric Lander & David Sabatini (Wang et al., 2014) as lentiviral particles, and colonies stably expressing Cas9 protein were selected using 2µg/ml Puromycin (Gibco). These stable Cas9-expressing cells were then transduced with viral particles carrying MLL sgRNA. After transduction, *mll* knock-out colonies were selected with 5µg/ml Blasticidin (Thermo Fischer Scientific). Several colonies were screened for loss of MLL protein expression through Western blot and finally, two clones (MLL iKO #11 and MLL iKO #20) were selected for further analysis. Cas9 expression (and therefore MLL knockout) was induced with 5µg/ml Doxycycline (Sigma) treatment for seven days where the medium was replenished after every three days. For CENP A-SNAP experiments, pcDNA FRT-CENP A-SNAP-3xHA was transfected into U-2OS and MLL mutant cells lines, and selected in media supplemented with 200μg/ml Hygromycin B. Similarly, the pcDNA 3.1-Puro-CENP A-SNAP-3xHA construct was transfected into U-2OS and SETD1A mutant cell lines, and selected using 4µg/ml Puromycin. Despite several attempts, we were unsuccessful in generating stable cell lines for pcDNA FRT-CENP A-SNAP-3xHA with MLLΔTAD. Therefore, pcDNA FRT-CENP-A-SNAP-3xHA vector was transiently transfected in MLLΔTAD cell line 12 hr. before siRNA transfection and assay was performed as depicted in Figure 6G.

### RNA interference

RNAi was performed with synthetic siRNA duplexes using Oligofectamine (Thermo Fischer Scientific) as previously described (Ali et al., 2017). The sequence of siRNA targeting the firefly luciferase gene (used as control) and various members of MLL family has been provided in supplemental information (Ali et al., 2014). Samples were collected at 48-72 hr after the first round of transfection as mentioned in the legends.

### Western blot

Whole-cell protein extracts were prepared by lysing cells in 2X NETN buffer (200mM NaCl, 40mM Tris-Cl pH 8.0, 1mM EDTA, and 1% NP-40) supplemented with a freshly prepared protease inhibitor cocktail (Sigma) and boiled for 10 min. Equal amounts of protein extracts were resolved by SDS-PAGE and transferred to either PVDF or nitrocellulose membrane. Immunoblotting was performed with the following antibodies: MLL (A300-374A, Bethyl Labs); SETD1A (A300-288A, Bethyl Labs); CENP-A (2186S, Cell Signaling Technology); CENP-B (ab25734, Abcam), CENP-C (ab50974, Abcam), HJURP (80508S, Cell Signaling Technology), HA (H6908, Sigma), and α-tubulin (T5168, Sigma). After probing with relevant secondary antibodies, blots were developed using Amersham™ ECL™ substrate or digital imaging using LI-COR Biosciences.

### Immunofluorescence

Cells (U-2OS, HEK-293, MLL, and SETD1A mutants expressing cell lines) used for immunofluorescence were grown on coverslips. Cells were arrested with nocodazole (100ng/ml) treatment for 12 to 16 hours and released into fresh medium for 60 mins (mitotic cells) or 90 mins (early G1), before fixing. Cells were fixed with freshly prepared 1% paraformaldehyde for 10 min at room temperature followed by permeabilized with 0.2% Triton X-100/PBS for 10 min. For staining RNA-DNA hybrids, cells were fixed and permeabilized with 100% ice-cold methanol for 10 min followed by acetone for 1 min on ice. Following fixation, immunofluorescence staining protocol was followed as described earlier (Ali et al., 2017). The samples were mounted in VECTASHIELD Mounting Medium (Vector laboratories-H1200) with 4’-6-diamidino-2-phenylindole (DAPI) to stain the DNA. Images were taken using a ZEISS LSM LSM 700 inverted confocal microscope with a 63x/1.4 oil immersion and quantified with either ZEN (ZEISS Efficient Navigation) or Image J software. Primary and secondary antibodies used for immunofluorescence were as follows: CENP-A (ab13939, Abcam), CENP-B (ab25734, Abcam), CENP-C (ab50974, Abcam), HJURP (80508S, Cell Signaling Technology), MLL_C_ (A300-374A, Bethyl Labs); MLL_N_ (A300-086A, Bethyl Labs); SETD1A (A300-288A, Bethyl Labs), S9.6 (MABE1095, Sigma or ENH001, Kerafast), total RNA Polymerase II (sc-9001, Santa Cruz Technology), and RNA polymerase II^S2P^ (ab24758, ab252855, Abcam), Alexa Fluor 488 (A11029, A11034, Invitrogen), Alexa Fluor 594 (A11032, A11037, A21209, Invitrogen).

For IF signal intensity quantification, Z-stack images with 0.5µm step size were taken. To quantify centromeric signal of CENP-A, CENP-B, CENP-C, and HJURP, the signal intensity was calculated by manually placing a circle (of equal radius) around the centromere in maximum intensity projection images and the pixel value of each channel was calculated. For each channel the background intensity was also calculated by placing another circle adjacent to the centromere signals, which was then subtracted from the respective IF channel value. To quantify co-localization of MLL and SETD1A with CENP-A in mitosis and early G, single plane images were used. Pearson correlation coefficient analysis was perform using ZEN black software.

### Chromatin immunoprecipitation

Chromatin immunoprecipitation (ChIP) was performed as described previously (Zargar et al., 2018, Sridhara et al., 2017). Briefly, ∼80% confluent HEK-293, IMR-90 tert, MLL inducible knock-out cells were fixed with 1% formaldehyde for 10 min at room temperature to perform cross-linking and quenched with 250 mM glycine for 5 min (also known as X-ChIP). Cells were lysed and sonicated to shear chromatin to achieve ∼200-500bp fragments. However, to immunoprecipitate centromeric proteins (CENP-A and CENP-C), native, non-crosslinked chromatin immunoprecipitation (also known as N-ChIP) was performed using a modified protocol described earlier (Carvalho et al., 2014). Briefly, cells were biochemically fractioned to get whole nuclei (de Almeida et al., 2010). To fragment the chromatin, whole nuclei were resuspended in 100ul MNase digestion buffer (15mM NaCl, 10mM Tris-Cl pH 7.4, 60mM KCl, and 1mM CaCl_2_) with 4U of MNase (N3755, Sigma) incubated at 37° C for 10 min. The reaction was inhibited immediately by quick-chilling on ice and by the addition of 100μl MNase stop buffer (100mM EDTA, 100mM EGTA, 0.05% NP-40). Then ChIP lysis buffer was added and incubated on ice for 15-30min.

For both kinds of ChIP experiments, 1/10^th^ of the fragmented chromatin was taken aside as input. The following antibodies were used for ChIP experiments: H3K4me2 (ab32356, Abcam); H3K36me2 (ab9049, Abcam); H3K9me3 (ab8898, Abcam); H3K9Ac (ab4441, Abcam), MLL (A300-374A, Bethyl Labs), SETD1A (A300-288A, Bethyl Labs), total RNA polymerase II (14958, Cell Signaling Technology or sc-9001, Santa Cruz Technology); RNA polymerase II^S2P^ (ab252855, Abcam); CENP-A (ab13939, Abcam), CENP-B (ab25734, Abcam), CENP-C (ab50974, Abcam), and IgG (12-370, Sigma). After incubation with primary antibodies overnight at 4°C followed by the addition of Protein A or G Sepharose beads (GE Healthcare) for 2-3 hrs, immunoprecipitated material was washed with ChIP wash buffers. Immunoprecipitated and input DNA were subsequently purified using standard Phenol, chloroform, and isoamyl alcohol extraction. The relative occupancy or percent input of the immunoprecipitated protein at each DNA locus was estimated by RT-qPCR as follows: 100 X 2^(Ct Input – Ct IP),^ where Ct Input and Ct IP are mean threshold cycles of RT-qPCR on DNA samples from input and specific immunoprecipitations, respectively. To measure fold over control, fold change over the ChIP values obtained in the control cells was used. The primer sequences are listed in the supplemental information.

### DNA:RNA immunoprecipitation

DNA: RNA hybrids immunoprecipitation (DRIP) was performed as described earlier (Smolka et al., 2012, Sridhara et al., 2017) with the following modifications. Briefly, cells were lysed in 300ul lysis buffer (100 mM NaCl, 10 mM Tris pH 8.0, 25 mM EDTA pH 8.0, 0.5% SDS) and sonicated using Diagenode Bioruptor (5 cycles of the 30s ON and 30s OFF at low intensity) and incubated with 100µg/ml Proteinase K over-night at 37°C. Nucleic acids were extracted from phenol-chloroform extraction and resuspended in DNase/RNase-free water. Nucleic acids were fragmented using a restriction enzymes cocktail (50U each of EcoRI, BamHI, HindIII, and XhoI). Fragmented DNA served as inputs. About 2-5μg of fragmented DNA was digested with 40U RNaseH (New England Biolabs) for at least 24 hours at 37^0^C to serve as a negative control. After cleaning digested nucleic acids with phenol-chloroform extraction and re-suspended in DNase/RNase-free water, S9.6 antibody (MABE1095, Sigma) was added in a 1:1 ration of nucleic acid: antibody in binding buffer (10 mM NaPO_4_ pH 7.0, 140 mM NaCl, 0.05% Triton X-100) and incubated overnight at 4°C. Immunoprecipitated complexes were pull-down using Protein A Sepharose beads (GE Healthcare) were at 4^0^C for 2 hours. Isolated complexes were washed thrice with ice-cold binding buffer and once with TE buffer for 2 min each, before elution (50 mM Tris pH 8.0, 10 mM EDTA, 0.5% SDS, 5µg proteinase k) for 30 min at 55°C. Nucleic acids were extracted using standard procedures. The relative occupancy or percent input of the immunoprecipitated DNA-RNA hybrids at each locus was estimated by RT-qPCR as follows: 100 X 2 ^(CtInput− CtIP)^, where Ct Input and Ct IP are mean threshold cycles of RT-qPCR on samples from input and specific immunoprecipitations, respectively. To measure fold over control, fold change over the DRIP values obtained in the control cells was used. The primers sequences are listed in the supplemental information.

### RNA isolation and qRT-PCR

Total RNA isolation and cDNA preparation was made as described earlier (Ali et al., 2017). Briefly, total cellular RNA was isolated from MLL and SETD1A mutant cells or parent U-2OS cells using the TRIzol (Ambion, Inc.,) or using an RNA isolation kit (Zymo Research). To inhibit RNA polymerase II transcription, U-2OS cells were treated with the following inhibitors: 20 µM Triptolide (T3652, Sigma), 20 µM LDC000067 hydrochloride (SML2179, Sigma), and 20 µg α-amanitin (A2263, Sigma) for 4 hrs. For cDNA synthesis, the isolated RNA was treated with TURBO DNase (Thermo Fisher Scientific) for 30 min at 37°C to remove genomic DNA contamination and no-enzyme RNA amplification was done before cDNA synthesis to ensure that purified RNA did not have DNA contamination. cDNA was synthesised using SuperScript™ III Reverse Transcriptase kit (Thermo Fischer Scientific) according to the manufacturer’s protocol. RT-qPCR was performed in either 7500 Real-Time PCR (Applied Biosystems), QuantStudio™ 5 Real-Time PCR System (Applied bioscience) or Bio-Rad (CFX-maestro) using DyNAmo ColorFlash SYBR Green qPCR kit (Thermo Fisher Scientific). The transcript levels were quantified using 2^-ΔΔ Ct^ (Livak and Schmittgen, 2001). The primers sequences are listed in the supplemental information.

### SNAP quench-pulse labelling of nascent CENPA

SNAP quench-pulse labelling was performed as described earlier (Jansen et al., 2007) with the following modifications. Parent U-2OS, MLL, and SETD1A mutant cells stably expressing CENPA-SNAP-HA were seeded on coverslips and treated with Control, MLL, or SETD1A siRNA for 48 hrs. Cells were arrested with thymidine (2µM) for 12 hrs followed by the treatment of 5 μM O6 –BG (BG-block) in complete growth media for 30 min at 37°C to quench the SNAP activity. The Blocker was removed by washing cells twice with PBS, once with medium, and finally replenished with a fresh growth medium to ensure the complete removal of a blocker reagent. Cells were then arrested with nocodazole (100ng/ml) for 12 hrs and released for 3hrs followed by treatment of TMR-Star (2μM, Covalys) for 30 min and stained using HA antibody (H6908, Sigma).

### Quantification and Statistical Analysis

GraphPad Prism 9.3 software was used to perform statistical analysis. Student t-test, and two-way ANOVA were performed as mentioned in the legends. In two-way ANOVA, significance is calculated against mean of control vs mean of test (or IgG). Error bars represent Standard Error Mean (SEM) or Standard Deviation (SD) wherever mentioned in the legends. For exact number of cells and experiments, please refer to figure legends.

## Supporting information

none

## Author contribution

K.K.M. made stable cell lines and performed experiments presented in Figures 1, 2A-B, E-F, 4D-E, 5A,C,E, 6A-F, SF1, SF2A-E,J, and SF4C-H. K.A.L. made MLL iKO cell lines and performed experiments presented in Figures 2C-D, G-H, 3B-C, and SF2 F-I, SF3 A-B and SF5A. S.C.S performed experiments related to R-loop and those presented in Figures 3A, D-F, 4A-C, 6K-L, SF3 C-E, SF4 A-B and SF5 B-D. P.D.K. made SNAP-CENP-A cell lines and performed experiments presented in Figures 5B,D,F 6G-J, and SF5E-H. K.K.M., S.C.S, P.D.K., K.A.L, and S.T, designed and analyzed the experiments. K.K.M., S.C.S, P.D.K and S.T. wrote the manuscript.

## Acknowledgments

We thank R. Roeder, F. Zhang, L. Jansen, S. Huang, E. Lander, and D. Sabatini for cDNA constructs. We would like to thank A. Karole for cloning MLL and SET1A shRNA and sgRNA constructs, J. Thakur for her help with initial experiments, and S.E.F facility for technical support. We also thank I. Cheeseman, A. Desai, M.D. Blower and W.C. Earnshaw for suggestions and discussions. K.K.M is recipient of Junior and Senior Research Fellowships of the University Grants Commission (UGC), India toward the pursuit of a Ph.D. degree of the Manipal University. K.A.L. and P.D.K. are recipients of Junior and Senior Research Fellowships of the Council of Scientific and Industrial Research (CSIR), India toward the pursuit of a Ph.D. degree of the Regional Centre for Biotechnology (RCB) and Manipal University respectively. This work was supported in part by the DBT/Wellcome Trust India Alliance Senior Fellowship to S.T.[IA/S/18/2/503981] and CDFD core funds.

