## Supplementary material for "MLL family members regulate H3K4 methylation to ensure CENP-A assembly at human centromeres": none

### Supplemental information Malik et al., 2022

#### Materials and Methods:

##### shRNA cloning and transfections:

shRNAs constructs for MLL and SETD1A were generated by cloning oligonucleotides (sequences given below) in EcoR1 and Kpn1 linearized pLKO.1 vector (Sigma). SETD1A shRNA#2 was a gift from Suming Huang (Deng et al., 2013). To deplete MLL and SETD1A using shRNA, HEK 293 cells were transiently transfected twice with MLL shRNA #1 and # 2 or SETD1A shRNA #1 and # 2 using PEI. Scrambled shRNA was used as control. Samples were collected 72 hr after the first round of transfection.

**shRNA sequences used in this study**:

| Name | Sequence | Source reference |
|---|---|---|
| MLL shRNA#1 | 5'-GCCTCCATCAACAGAAAGGAT | This paper |
| MLL shRNA #2 | 5'- CTACCAACCCTAAACCCTG | Liu et al., 2007 |
| SETD1A shRNA #1 | 5'-AGCAAAAGGGACCCACCCC | This study |
| SETD1A shRNA #2 | 5'GACAACAACGAATGAAATA | Deng et al., 2013 |
| Scrambled | 5'-GCGCGATAGCGCTAATAATTT | Liu et al., 2007 |

**siRNA sequences used in this study**:

| Name | Sequence | Source reference |
|---|---|---|
| MLL siRNA #1 | 5'-AAGGAAAGCAUUACUGAGAAAUU | Ali et al., 2014 |
| MLL siRNA #2 | 5'-ACGAAAGACTGAATGTAAAUU | Ali et al., 2014 |
| SETD1A siRNA$^{SR}$ #1 | 5´-GGAAAGAGCCAUCGGAAAUUU | Ali et al., 2014 |
| SETD1A siRNA #2 | 5´-GCAAGAUGGUGGAGAACGUUU | Ali et al., 2014 |
| SETD1B siRNA | 5'-GAUGAGAACCAAUGAGUUUUU | This paper |
| MLL2 siRNA #2 | 5´-GGAUGAAGUUAGAGAAAAUUU | Ali et al., 2014 |
| MLL3 siRNA #2 | 5´-GGAUAGAGCUAAGGGAUAAUU | Ali et al., 2014 |
| Luciferase siRNA | 5´-CGUCGCGGAAUACUUCGA | Ali et al., 2014 |

**The sgRNAs used for MLL are as follows**:

| Name | Primer designation | Sequence |
|---|---|---|
| MLL sgRNA 1 | Fw | CACCGACATGGCGCACAGCTGTCGG |
| | Rv | AAACCCGACAGCTGTGCGCCATGTC |
| MLL sgRNA 2 | Fw | CACCGCGAACATGGCGCACAGCTGT |
| | Rv | AAACACAGCTGTGCGCCATGTTCGC |

**The sequence of primers used in this study for ChIP, DRIP, and transcript analysis:**

| Gene | Primer designation | Sequence | Application | Source or reference |
|---|---|---|---|---|
| *α-satellite* | Fw | CATCACAAAGAAGTTTCTGAGAATGCTTC | ChIP, DRIP, and transcript analysis | Quénet et al., 2014 |
| | Rv | TGCATTCAACTCACAGAGTTGAACCTTCC | | |
| *D17Z1* | Fw | CTTTGGATGGAGCAGGTTTGAGAC | ChIP, DRIP, and transcript analysis | McNulty et al., 2017 |
| | Rv | CGTTTAGTTAGGTGCAGTTATCC | | |
| *D17Z1-B* | Fw | CACTGTTTGGCCTTCGTTC | ChIP, DRIP, and transcript analysis | McNulty et al., 2017 |
| | Rv | TCCACTTGCAGATTCCACA | | |
| *D17Z1-C* | Fw | GCCTATGGTACTAAAGGGAAT | ChIP, DRIP, and transcript analysis | This study |
| | Rv | ATCCTCAGAGAGGTCCAAAT | | |
| *HOXA9* | Fw | GTGGACTCGTTCCT | Transcript analysis | This study |
| | Rv | CTTGGACTGGAAGCTGCA | | |
| *HOXA9* | Fw | CTCCGCCGCTCTCATTCTCAG | ChIP | This study |
| | Rv | GCCAGAAGGGGTGACTGTCC | | |
| *PAX3* | Fw | AGCCGCATCCTGAGAAGTAA | Transcript analysis | This study |
| | Rv | CAGCTGTTCTGCTGTGAAGG | | |
| *PAX9* | Fw | CCGCATGACAGATTTTGCTA | ChIP | This study |
| | Rv | GCGTTTGGTCTGAATGTGAA | | |
| *RAD18* | Fw | ATGCGCAGTACAAGCCCTTA | ChIP | This study |
| | Rv | GCTCCAACACCACTCGAAAT | | |
| *RAD18* | Fw | GGATGCGTCTTGAAGCTAGTAA | Transcript analysis | This study |
| | Rv | CTGATCCACCAGAAGCTGAAA | | |
| *RPL13A (Intron 7)* | Fw | AGGTGCCTTGCTCACAGAGT | DRIP | Sanz et al., 2019 |
| | Rv | GGTTGCATTGCCCTCATTAC | | |

| | | | | |
|---|---|---|---|---|
| *RPL13A (Exon 8)* | Fw | GAGCAAGGAAAGGGTCTTAG | DRIP | This study |
| | Rv | CTTCTAGAAATACCCTGTGTAC | | |
| *MLL* | Fw | GGAGCACACATTCCAGACCA | Transcript analysis | Ali et al., 2014 |
| | Rv | TTTGGGTCACCTGAACTTCC | | |
| *MLL2* | Fw | AGCCGTGTGAGGATGAAAAC | Transcript analysis | Ali et al., 2014 |
| | Rv | ACCTGGGGAGGACCATCTT | | |
| *MLL3* | Fw | CAGCACCACGAAAACAAAGA | Transcript analysis | This study |
| | Rv | ACTCCACACAACGGTGATGA | | |
| *SETD1A* | Fw | CGAATACGTGGGTCAGAACA | Transcript analysis | Ali et al., 2014 |
| | Rv | TGCAGCAGTGGTTGATGAAT | | |
| *SETD1B* | Fw | TGGACACCAAAGGGGAAACC | Transcript analysis | This study |
| | Rv | CAGACAGGCCTCCATCCTTG | | |
| *GAPDH* | Fw | CGAGATCCCTCCAAAATCAA | Transcript analysis | This study |
| | Rv | TTCACACCCATGACGAACAT | | |
| $U2_C$ | Fw | TTTGCTCCCACTGCCGTC | ChIP and DRIP | Zargar et al., 2018 |
| | Rv | CTGAGTCTTTCGGTGCCC | | |
| *CD4* | Fw | TCTGCAGAAGGAACAAAGCA | ChIP | This study |
| | Rv | GGAAGGAAGCCGAGTCTGA | | |
| *EGR1* | Fw | GAACGTTCAGCCTCGTTCTC | DRIP | Sanz et.al., 2016 |
| | Rv | GGAAGGTGGAAGGAAACACA | | |
| *SNRPN* | Fw | TGCCAGGAAGCCAAATGAGT | DRIP | Sridhara et al., 2017 |
| | Rv | TCCCTCTTGGCAACATCCA | | |

#### Supplemental Figure Legends

**Figure S1:RNAi-mediated downregulation of MLL family members abrogates centromeric transcription**.

(**A**) siRNA mediated downregulation of various MLLs was performed and the efficacy of siRNA treatment was determined by plotting the qRT-PCR analysis of respective transcript levels after total RNA extraction and cDNA synthesis. (**B**) cDNA samples obtained after RNAi treatment of Control, MLL2, MLL3, and SETD1B (from **A**) were analysed for cenRNA transcripts from D17Z1, D17Z1-B, and D17Z1-C α-satellite arrays of chromosome 17. (**C**) qRT-PCR analysis of α-satellite cenRNA from individual HOR of chromosome 17 and RNA Pol II regulated gene *PAX3* after treatment with either control (DMSO), Triptolide (20 µM), CDK9 inhibitor (20 µM), α-amanitin (20 µg) for 4 hr. is shown. (**A-C**) cDNA was synthesized from total RNA after rigorous DNase I treatment and amplified using qRT-PCR for indicated RNAs. Data from all samples were normalized to GAPDH mRNA levels from respective samples by using − ΔΔCT method and expression is shown relative to control siRNA-treated/DMSO treated cells from respective cell line/treatment (which is arbitrarily set to 1). Each experiment was performed at least three, or more times except α-amanitin treatment (two times). Error bars represent SD. *P ≤ 0.05, **P ≤ 0.005, *** P ≤ 0.0005, ****P ≤ 0.0001, ns: not significant P > 0.05 (two tailed Student's t-test). (**D-E**) Immunoblots of whole-cell lysate were prepared from cells treated with either control and MLL (**D**) or SETD1A siRNA (**E**). Blots were probed with α-MLL$_C$ (**D**) / α-SETD1A (**E**) and α-tubulin as shown. Molecular weight markers (in kDa) are shown on the left. (**F**) Schematic representation of recombinant MLL and SETD1A mutants used in this study displaying different domains in these proteins. Full-length MLL (FL), and SET domain deleted MLL (Δaa3829–3969) U-2OS cell lines have been described before (Ali et al., 2014). MLLΔTAD was generated in full-length MLL here by deletion of aa 2847–2855 using site-directed mutagenesis. U-2OS cells expressing full-length SETD1A (FL), SET domain deleted SETD1A (Δaa1407-1707) or point mutation inactivating SET domain (SETD1A N1646A mutant) were generated from siRNA resistant full-length SETD1A cDNA described in **G**. (**G**) siRNA resistant full-length SETD1A was generated by introducing seven silent mutations (shown in red) at wobble positions between nucleotide +2916 to +2934 (Accession no NM_014712.3) in the full-length construct using site-directed mutagenesis. The siRNA (siRNA#1) sequence is underlined. CDK9i, CDK9 inhibitor; AT-hook: AT-rich region; F, FLAG epitope tag; PHD, Plant homeodomain; Zn CXXC: Zinc-finger domain; Bromo: Bromodomain; RRM, RNA recognition motif; FYRN/C: Phenylalanine and tyrosine-rich region N-terminal/C-terminal; Win, WDR5 interacting motif.

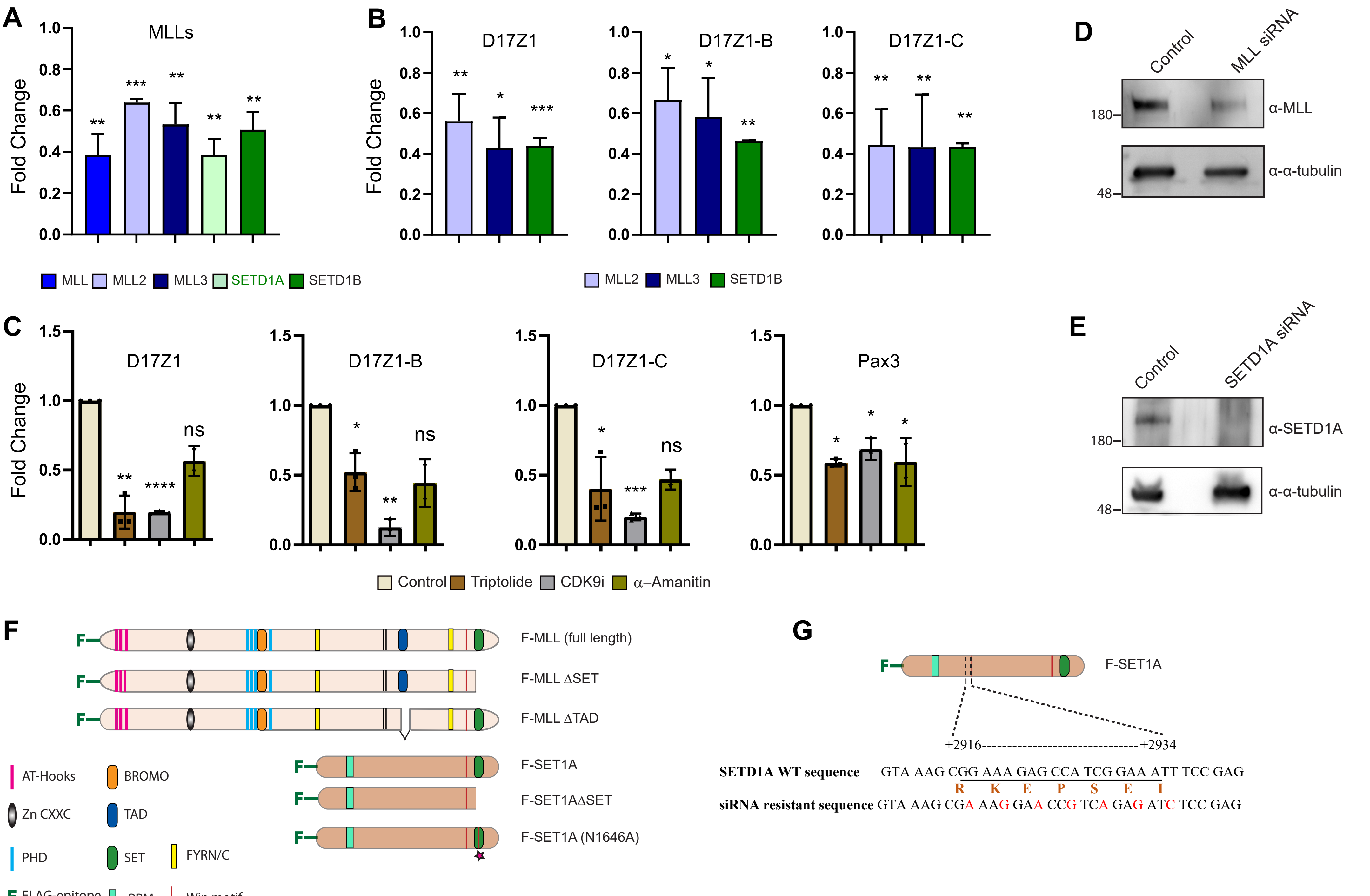


Malik Supplementary Figure 1

**Figure S2: MLLs bind to the human centromere repeats.**

(**A**) Immunofluorescence staining (IF) of endogenous MLL (green) or SETD1A (green) with CENP-A (red) in U-2OS cells in interphase is shown. DNA was stained with DAPI (blue). The area in the white square is magnified and shown on the right for each image. (**B**-**C**) U-2OS cells were transfected with siRNA specific to Control, MLL (**B**), or SETD1A (**C**) to check for the specificity of MLL or SETD1A staining at the centromere. Cells were stained with endogenous MLL (green) or SETD1A (green) with CENP-C (red) or CENP-A (red), and DNA was stained with DAPI (blue) as indicated. (D) U-2OS cells, stained with Alexa Flour 488 and Alexa Flour 594, are shown. (**A**-**D**) Scale bar, 5μm. (**E**) U-2OS cells were stained with MLL2 (green), MLL3 (green), or SETD1B(green) antibody along with centromeric marker CENP-A (red) as shown. The area in the white square is magnified and shown on the left for each image. Scale bar, 2μm. (**F, H**) Immunoblot show MLL (**F**) and SETD1A (**H**) shRNA (#1 and #2) knockdown efficiency in treated HEK-293 cells. The blots were probed with α-MLL (**F**) or α-SETD1A (**H**), and α-α-tubulin antibody. (**G, I**) Chromatin immunoprecipitation (ChIP) analyses showing enrichment of MLL (**G**) and SETD1A **(I)** at centromeric α-satellite loci following treatment of either MLL shRNA or SETD1A shRNA in HEK-293 cells, and the result plotted as percent input enrichment, are shown. Each experiment was performed at least three, or more times. Error bars represent SD. $^{*}P \leq 0.05$, $^{**}P \leq 0.005$, $^{***} P \leq 0.0005$, ns: not significant $P > 0.05$ (two tailed Student's t-test). (**J**) CENP-B occupancy, detected by ChIP, is shown. Data from three or more independent ChIP experiments is shown. Error bars represent SD. $^{*}P \leq 0.05$, $^{**}P \leq 0.005$, ns: not significant $P > 0.05$ (Two-way ANOVA with Šídák multiple comparison test). α-sat, α-satellite.

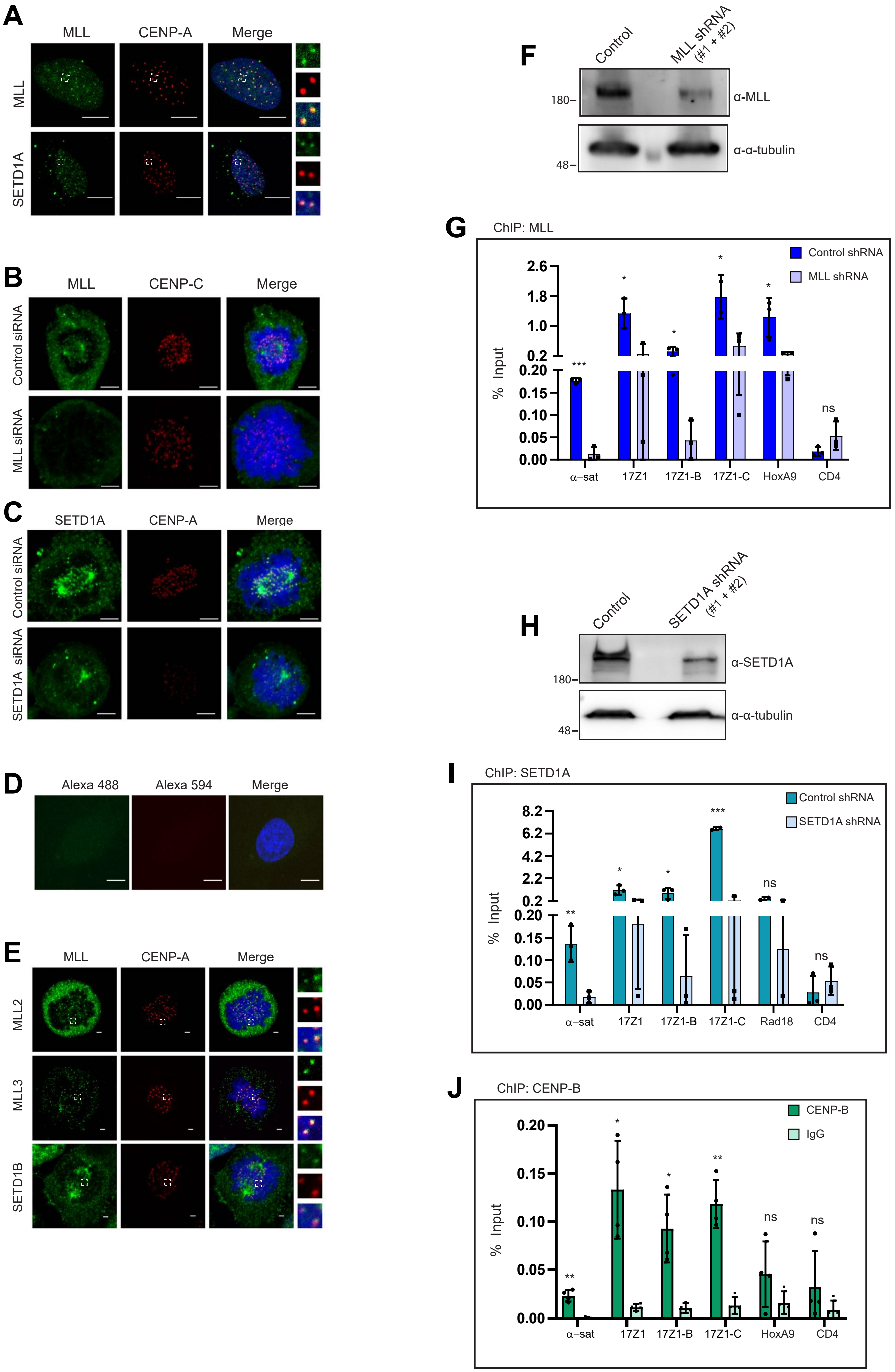


Malik Supplementary Figure 2

**Figure S3: Loss of MLL affects the epigenetic landscape of the centromeres.**

(**A**-**E**) ChIP-analyses with MLL (**A**), H3K4me2 (**B**), H3K9ac(**C**), H3K9me3 (**D**), and H3K36me2 (**E**) antibodies in *MLL* iKO cells (#20) is shown. Data were normalized against the ChIP values obtained in parental (or Cas9-expressing) cells, which are used as Control. Data from three or more independent ChIP experiments are plotted. Error bars represent SD. *P ≤ 0.05, **P ≤ 0.005, *** P ≤ 0.0005, ****P ≤ 0.0001, ns: not significant P > 0.05 (Two-way ANOVA with Šídák multiple comparison test). α-sat, α-satellite.

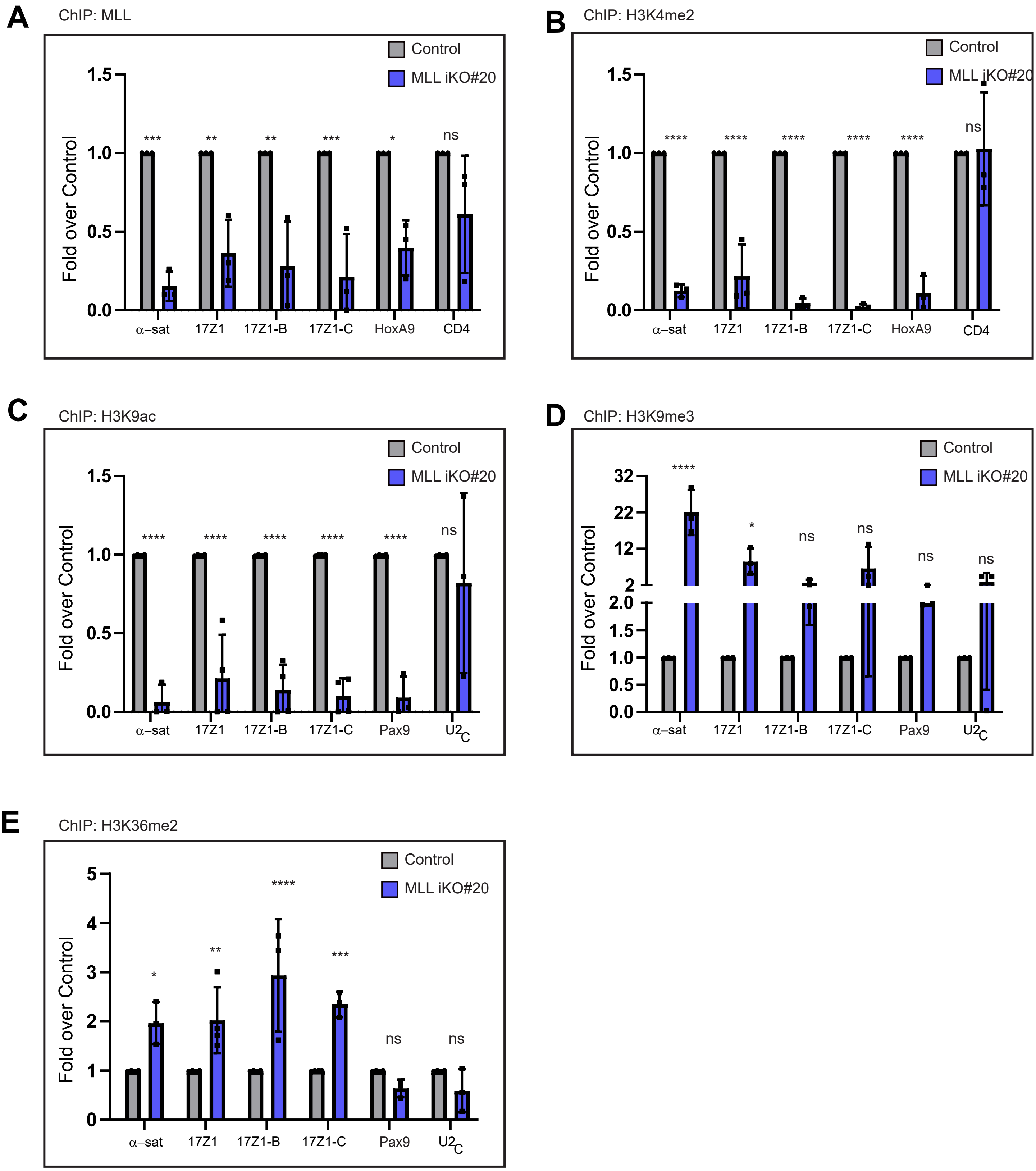


Malik Supplementary Figure 3

**Figure S4: Disparate impact of MLL and SETD1A on centromeric R-loops.**

(**A**) Representative images for the data represented in Figure 4A show nuclear R-loops, 48 hrs after MLL and SETD1A siRNA treatment in U-2OS cells. The cells were stained using the S9.6 (green) antibody and DAPI (blue). Each nucleus is outlined in white. For transcription inhibition, cells were treated with 20 µM Triptolide or DMSO (Control) for 4 hrs. Scale bar, 10 µm. (**B**) DRIP analysis in MLL RNAi-depleted HEK-293 cells, with respective RNase H controls, is shown. Data are presented as percent input enrichment. Data from three or more independent DRIP experiments are plotted. Error bars represent SD. **P ≤ 0.005, ns: not significant P > 0.05 (Two-way ANOVA with Šídák multiple comparison test). Ctrl, control; si, siRNA. **(C-D)** Immunofluorescence staining of Total RNA Pol II (**C**, green) or RNA Pol $II^{S2P}$(**D**, green) and CENP-A (red) in mitotic cells following treatment with either Control, MLL, or SETD1A siRNA, are shown. Scale bar, 5µm. (**E-F**) Quantification of fluorescence intensity at the centromere of Total RNA Pol II (**E**) or RNA Pol $II^{S2P}$(**F**) in Control, MLL, or SETD1A siRNA treated cells as shown in **C-D**. Each data point represents a single centromere. Maximum Intensity projection (MIPs) images were used in all quantification analyses using ZEN software. n ≥ 300 centromeres, (n=2 experiments). ****P ≤ 0.0001, ns: not significant p > 0.05; (Mann-Whitney two-tailed unpaired test). (**G-H**) ChIP-analysis of RNA Pol II (**G**) and RNA Pol $II^{S2P}$ (**H**) in *MLL* iKO #11 cells are shown. Data were normalized against the ChIP values obtained in parental (or Cas9-expressing) cells, which are used as Control. Data from two independent ChIP experiments are plotted. Error bars represent SD. *P ≤ 0.05, **P ≤ 0.005, ****P ≤ 0.0001, ns: not significant P > 0.05 (Two-way ANOVA with Šídák multiple comparison test). Ctrl, control; si, siRNA.

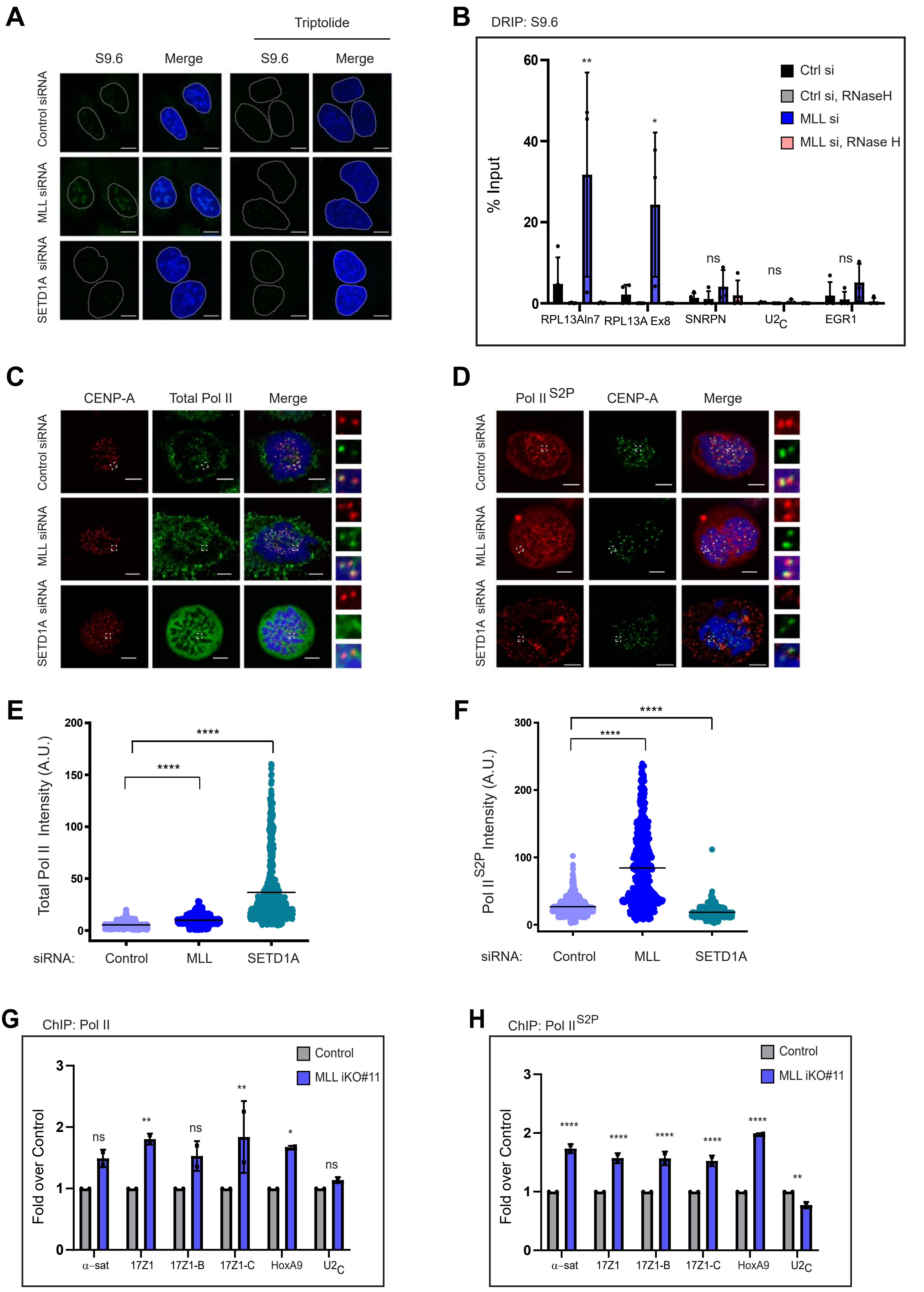


Malik Supplementary Figure 4

**Figure S5: MLL and SETD1A facilitate recruitment of nascent CENPA at centromeres.** (**A**-**D**) Immunoblot shows CENP-C (**A**), CENP-B (**B**), HJURP (**C**), and CENP-A (**D**) protein levels in inducible *MLL* knock outs (iKO #11 and #20) cells. Blots were probed with respective antibodies as indicated. Molecular weight markers (in kDa) are shown on the left. (**E**) Quantification of centromeric fluorescence intensity of ectopically expressed total CENP-A (stained using α-HA antibody) in parent U-2OS cells (—) or cell line stably expressing MLL full length (FL) or MLLΔTAD or MLLΔSET upon MLL siRNA treatment. ****$P \leq 0.001$, ns: not significant $P > 0.05$ (Mann-Whitney two-tailed unpaired test). (**F**) Quantification of centromeric fluorescence intensity of total CENP-A in parent U-2OS cells (—) or cell line stably expressing siRNA resistant SETD1A full length (FL) or SETD1AΔSET (here N1646A mutant was used) upon SETD1A siRNA. ****$P \leq 0.001$, ns: not significant $P > 0.05$ (Mann-Whitney two-tailed unpaired test). (**E-F**) Representative images are shown in Figure 6H. Each data point represents a single centromere. $n \geq 300$ quantified from 10 early G1 cell pairs, (n=2 experiments). Maximum Intensity projection (MIPs) images were used in all quantification analyses using ZEN software. (**G**-**H**) Stably expressing CENP-A SNAP-3xHA cells were treated with MLL (**G**) or SETD1A (**H**) and Control siRNA, and collected after 72 hrs for cell lysate preparation. CENP-A-SNAP-3xHA protein was detected using an α-HA antibody. A.U., arbitrary units.

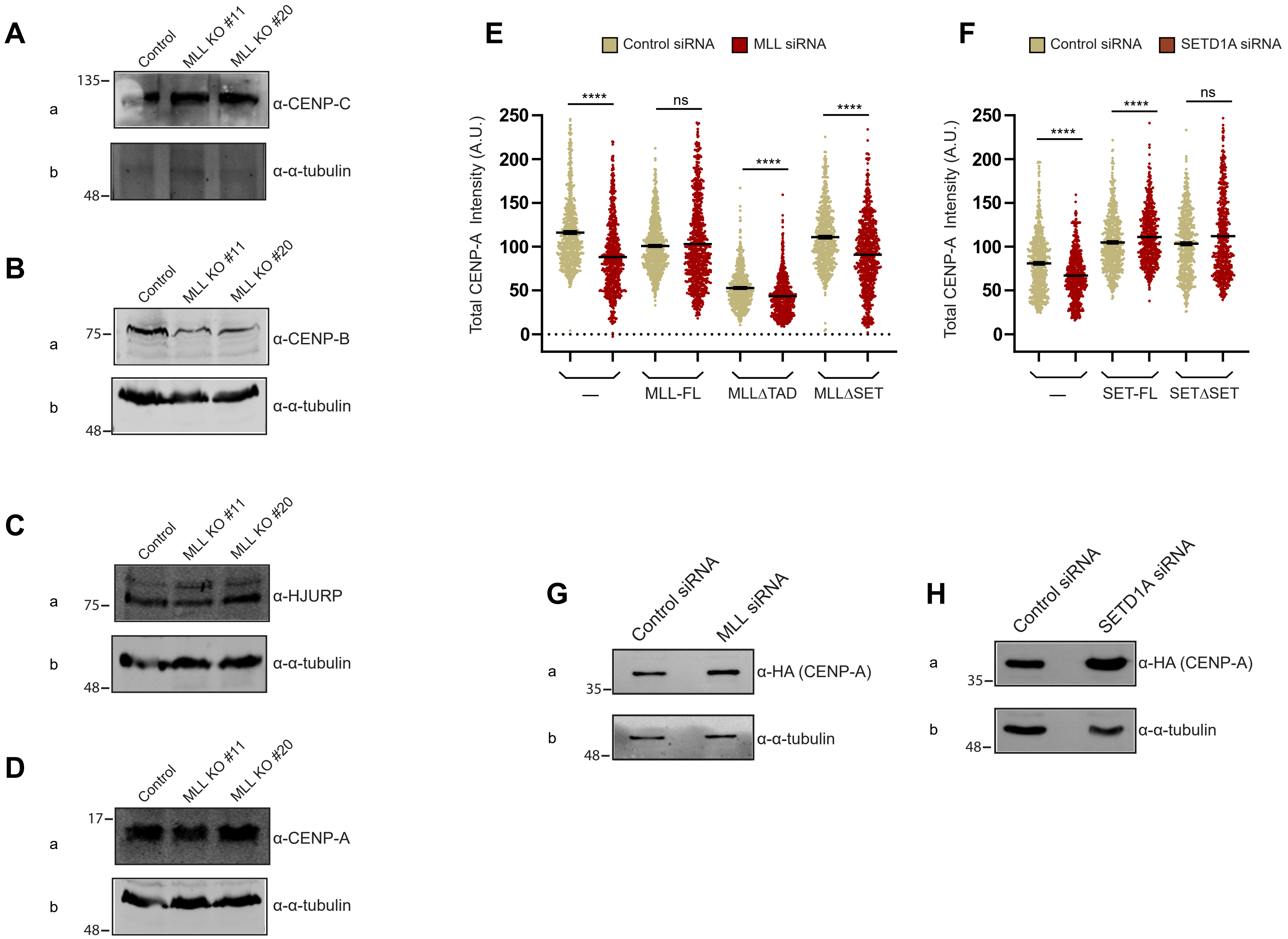


Malik Supplementary Figure 5
